# Interpretable Prediction of Phase Separation and Disease Variant Effects in Intrinsically Disordered Regions

**DOI:** 10.64898/2026.07.14.736885

**Authors:** Mingjie Zhao, Sushant Kumar

## Abstract

Coding mutations within intrinsically disordered regions (IDRs) of proteins are increasingly implicated in human diseases yet remain poorly interpreted by conventional variant-effect predictors that rely on structural stability and conservation-based metrics. Quantifying disruption of IDR-mediated liquid-liquid phase separation (LLPS) offers a biophysically principled approach to interpreting the pathogenic impact of such variants. However, existing LLPS predictors suffer from training biases toward self-separating proteins, show limited performance on partner-dependent phase separation, and often lack interpretability for variant prioritization. We present an interpretable ensemble machine-learning framework that integrates protein language model embeddings of sequence and predicted structure to predict LLPS propensity and classify proteins as self-separating or partner-dependent. Our two-step classifiers outperform existing methods on independent benchmark datasets, with the largest gains for partner-dependent LLPS proteins. Beyond classification, our framework identifies critical phase-separating regions and quantifies mutation-induced perturbations in LLPS. Applied to disease-associated variant databases, we found that pathogenic mutations are enriched in predicted phase-separating regions and frequently perturb LLPS propensity scores, implicating mutation-induced LLPS dysregulation as a potential pathogenic mechanism for numerous diseases. Overall, our framework provides an accurate, interpretable approach for identifying phase-separating proteins and linking aberrant phase-separation behavior to disease pathogenesis.

## Introduction

Intrinsically disordered regions (IDRs) constitute approximately one-third of the eukaryotic proteome^1^. IDRs frequently mediate interactions with other proteins, nucleic acids, and small molecules, thereby promoting the formation of higher-order molecular assemblies^2^. These IDR-mediated interactions and associated higher-order molecular assemblies are essential for biological processes, including cellular signaling^3,4^, DNA damage repair^5^, transcription^6^, and metabolism^7^. Therefore, disruption of these IDR-mediated interactions has increasingly been implicated in various diseases, including neurodegenerative disorders, cancers, and a range of rare diseases^8–12^. Prior studies have shown that a substantial proportion of disease-associated coding variants, particularly missense substitutions, map to these disordered regions^10^. However, the functional consequences of such variants remain difficult to interpret with current variant effect predictors (VEPs). Most VEPs primarily leverage protein structure or intra- and inter-species evolutionary constraints to quantify pathogenicity scores based on loss of protein structural stability, disruption of binding interfaces, or destabilization of the native fold^13–15^. Furthermore, these predictors are often benchmarked primarily against variants within folded domains of proteins. In contrast, IDRs lack well-defined three-dimensional structure and exhibit heterogeneous conformational ensembles. Therefore, interpreting missense mutations within IDRs using a structure-centric view poses a significant challenge^10^, leaving a mechanistically important and potentially clinically relevant class of mutations relatively underexplored in systematic variant-interpretation efforts. This knowledge gap necessitates the development of novel approaches that primarily rely on the biophysical principles underlying IDR function, rather than on structural stability, to guide pathogenicity prediction for mutations affecting these regions.

One approach to quantify the molecular impact of genetic variants on IDRs is to assess disruption of IDR-mediated liquid-liquid phase separation (LLPS). LLPS is a biochemical process in which proteins and nucleic acids spontaneously demix from a homogeneous solution to form liquid droplets. These demixing events are driven primarily by weak, multivalent interactions among aromatic, charged, and other “sticker” residues, which are often found within IDRs^16,17^. LLPS enables intracellular compartmentalization and underlies the formation of membrane-less organelles (also known as biomolecular condensates), including stress granules, P-bodies, the nucleolus, transcriptional condensates at super-enhancers, DNA damage foci, and signaling clusters at the plasma membrane^18–20^. Proteins that undergo LLPS can be further classified as self- separating (PS-Self) or partner-dependent (PS-Part) based on their mode of phase separation^21^. For instance, PS-Self proteins employ homotypic, multivalent interactions within their disordered regions to drive self-assembly, as exemplified by FUS, TDP-43, and hnRNPA1 proteins^22,23^. In contrast, PS-Part proteins rely on heterotypic interactions with other biomolecules (proteins, RNA, and chromatin) to drive higher-order assemblies^24^.

Recent studies increasingly recognize that genetic alterations promoting aberrant LLPS may serve as a pathogenic mechanism across various human diseases^25^. For instance, missense mutations in the IDRs of FUS, TDP-43, and hnRNPA1 have been shown to alter the functions of associated stress granules and related condensates, leading to amyotrophic lateral sclerosis and frontotemporal dementia^22,23^. Moreover, disease-associated mutations in tau proteins can accelerate aberrant phase transitions, leading to condensate dysregulation, which has been implicated in tauopathies^26^. Similarly, mutations in the tumor suppressor SPOP, the nucleolar protein NPM1, and a growing list of oncogenic fusion proteins, including EWS-LI1, have been shown to drive aberrant gain- or loss-of-condensates that contribute to tumorigenesis^19,27,28^. Despite these numerous examples, a systematic interpretation of variants affecting IDRs, measured by disruption of the LLPS properties of underlying proteins at scale, remains elusive.

Current experimental methods for detecting LLPS proteins primarily rely on immunofluorescence-based approaches, which are inherently limited by high costs, low throughput, and experimental confounders. Therefore, there has been growing interest in predicting LLPS proteins using feature engineering and data-driven machine learning methods. Feature-engineered machine learning methods predict LLPS propensity based on biophysical properties and sequence features, including weak multivalent interactions and the presence of IDRs, prion-like domains, and low-complexity regions^29–32^. While most of these methods produce a single LLPS score per protein, some tools, such as PhaSePred^30^, distinguish LLPS subtypes through separate PS-Self and PS-Part scores. More recent methods provide residue-level predictions by integrating sequence features with post-translational modifications and network properties^33^. Similarly, catGranule2 integrates sequence features with AlphaFold2-derived structural features to quantify residue-level LLPS propensity scores^34^. Notably, some of these methods enable in silico measurement of mutation-induced LLPS perturbations by comparing wild-type and mutant scores^33,34^. In parallel, methods leveraging protein language model (pLM) embeddings have shown significant improvements in LLPS prediction across benchmark studies^35–37^. Despite these advances, current approaches have several key limitations. For example, most predictors perform substantially better on self-separating proteins than on partner-dependent LLPS proteins, reflecting an inherent bias in training sets dominated by PS-Self proteins from various LLPS databases^38–40^. Similarly, data-driven and pLM-based methods typically operate as black boxes, and only a few approaches integrate sequence and structural representations into a unified framework for predicting LLPS proteins^41^. These gaps are particularly critical for variant interpretation, which requires accurate handling of both LLPS modes, residue-level attribution scores, and interpretability grounded in the biophysical features that drive phase separation.

We sought to address these gaps by developing an interpretable pLM-based framework to predict the LLPS propensity score from protein sequences and structures, and by characterizing how genetic variants affect these scores across various diseases. Our data-driven approach embeds raw protein sequences and predicted structures into condensed representations that capture context-dependent residue interactions, long-range dependencies, sequence motifs, and implicit biophysical patterns for predicting LLPS proteins. Specifically, we integrated sequence and structure embeddings within an ensemble neural network framework and trained two-step models. Our first classifier distinguishes LLPS from non-LLPS proteins, whereas the downstream classifiers further categorize LLPS proteins as PS-Self or PS-Part. Subsequently, we applied a sliding-window method to protein segments to quantify residue-level attribution scores and characterize physicochemical features, thereby identifying sub-regions that drive LLPS predictions and align with known phase-separating regions. By applying our models to sequence segments, we pinpointed critical regions that promote phase separation and were consistent with experiments. On independent validation and carefully curated benchmarking datasets, our framework outperformed existing methods in predicting LLPS, PS-Self, and PS-Part proteins, with the largest gains observed for the partner-dependent class. Finally, we applied this framework to disease-associated variant databases to identify many well-characterized and novel proteins in which pathogenic mutations either perturb LLPS or fall within predicted phase-separating regions. Overall, our study provides an accurate, interpretable framework for systematically evaluating how coding variants in IDRs may contribute to disease through aberrant phase separation.

## Results

### Overview of Our Computational Framework for LLPS Prediction and Mutation Evaluation

We developed an interpretable machine-learning framework that combines multimodal protein representations with a hierarchical classification strategy to systematically identify LLPS proteins. This approach enabled the identification of critical residues and the prediction of mutation-driven perturbations to phase-separation behavior. First, we generated embeddings capturing amino acid sequence information and predicted structural context using two pretrained protein language models, ESM2^36^ and SaProt^37^. We then trained multiple machine learning frameworks for LLPS prediction, including sequence-, structure-, and ensemble-based models. In the ensemble model, both modalities were integrated through attention-based mechanisms to produce a continuous LLPS propensity score (**Fig. 1A; Supplementary Fig. S1**). We systematically evaluated the performance of these models in predicting LLPS proteins. Subsequently, we implemented a hierarchical classification strategy to categorize LLPS proteins into self-assembling (PS-Self) and partner-dependent (PS-Part) subclasses (**Fig. 1B**). This hierarchical design enables the model to disentangle intrinsic LLPS capacity from mechanisms that rely on interaction partners.

**Figure 1.**
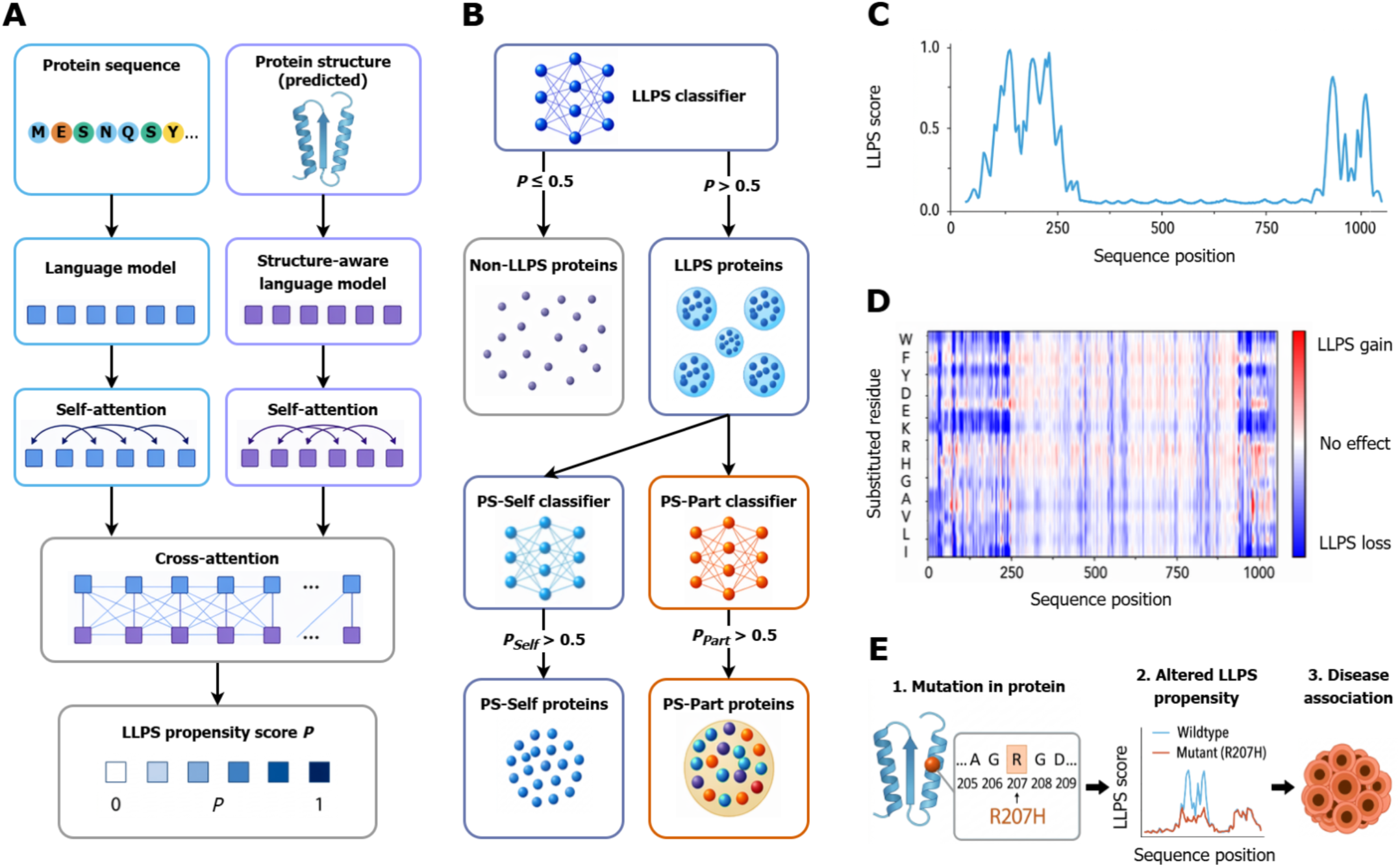
Overall schematic of our framework. **A)** An overview of the ensemble model architecture for LLPS protein classification. Protein sequences are embedded with a pretrained language model, ESM2, while structural information is embedded with a pretrained structure- aware language model, SaProt. The two embedding spaces are fused via a cross-attention layer. The output is an LLPS propensity score for each protein. **B)** Hierarchical classification framework for identifying LLPS proteins and distinguishing between PS-Self and PS-Part proteins. The ensemble model predicts whether a protein is likely to undergo LLPS. Proteins predicted to undergo LLPS are then passed to two downstream classifiers that independently predict PS-Self and PS-Part propensities. **C)** Residue-level LLPS-promoting regions identified by the ensemble model. **D)** Saturation mutagenesis analysis to evaluate the effects of missense mutations on phase- separation propensity. Differences between mutant and wild-type LLPS propensity scores are used to identify mutations predicted to enhance or disrupt LLPS behavior. **E)** LLPS dysregulation as a potential mechanism linking mutations to disease.

Our interpretable approach provides residue-level LLPS propensity profiles that reveal sequence features underlying the predictions. This enabled us to pinpoint the critical regions that contributed most to the LLPS propensity score, which frequently aligned with experimentally validated phase-separating regions (**Fig. 1C**). Additionally, we performed in silico saturation mutagenesis to systematically identify variants that either enhance or disrupt phase separation capacity (**Fig. 1D**). Finally, we integrated our predictions with several genetic variant databases, including ClinVar^38^ and the denovo-db^39^, to map LLPS-altering mutations to known pathogenic or *de novo* mutations, providing a mechanistic link between phase separation dysregulation and disease-associated mutations (**Fig. 1E**).

### Dataset description and model performance evaluation

We collected 4,857 LLPS proteins and split them into training and validation sets. A subset of these proteins (n=748) was drawn from a recent proteome-wide experimental study of endogenous condensate proteins^42^ and used as the training set. Additional LLPS proteins were obtained from the DrLLPS^40^, LLPSDB^39^, and PhaSePro^43^ databases, which categorize proteins by phase-separation role as PS-Self or PS-Part. From these databases, we gathered 352 PS-Self proteins (107 from DrLLPS, 230 from LLPSDB, and 15 from PhaSePro) and 3,757 PS-Part proteins (3,714 from DrLLPS, 31 from LLPSDB, and 12 from PhaSePro). We randomly sampled 116 PS-Self and 116 PS-Part proteins from this collection for the validation set, adding the remaining proteins to the training set. This resulted in 4,625 LLPS proteins in the training set and 232 LLPS proteins in the validation set (**Fig. 2A**). For non-LLPS proteins, we selected proteins from the PDB that show no current evidence of LLPS association, are not listed in any of the LLPS databases used here, and lack annotations of potential LLPS interactors. We randomly sampled 4,625 non-LLPS proteins for the training set and 232 for the validation set to maintain a balanced number of positive and negative samples (**Fig. 2A**). Additionally, we used an external dataset from an LLPS benchmark study^44^ to systematically evaluate and compare the performance of our models with other methods. This dataset includes annotations for PS-Self and PS-Part proteins and features both fully structured and intrinsically disordered non-LLPS proteins. After removing all proteins present in the training set, the final test set contains 285 PS-Self proteins, 222 PS-Part proteins, 42 proteins annotated as both PS-Self and PS-Part, 930 fully structured non-LLPS proteins, and 930 intrinsically disordered non-LLPS proteins (**Supplementary Table 1**).

**Figure 2:**
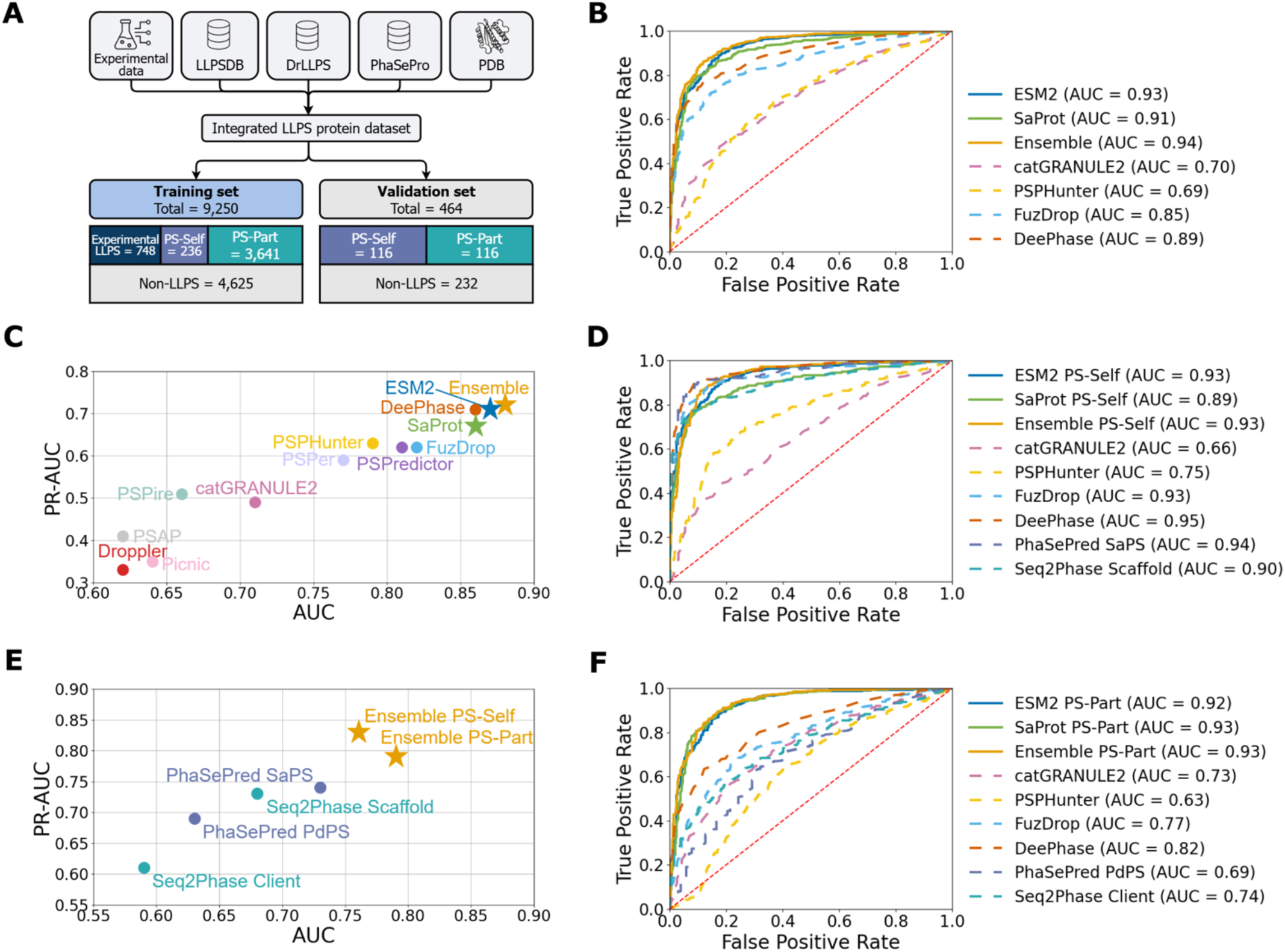
Model evaluation and benchmarking. **A)** Datasets used for model training and validation included experimentally identified LLPS proteins, annotated PS-Self and PS-Part proteins, and non-LLPS proteins. The integrated LLPS protein dataset was split into a training set (total = 9,250) and a validation set (total = 464). **B)** ROC curves on the validation set for LLPS vs. non-LLPS classification, comparing our ensemble model with models utilizing ESM2 and SaProt-based embeddings individually. ROC comparison with previously developed tools, including FuzDrop, DeePhase, catGRANULE2, and PSPHunter, is also presented. **C)** Area under the precision-recall curve (PR-AUC) versus area under the ROC curve (ROC AUC) for LLPS vs. non-LLPS classification on the independent test set, highlighting the superior performance of the ensemble model and ESM2. **D)** ROC curves on the validation set for PS-Self vs. non-LLPS classification, comparing the ensemble model with ESM2 and SaProt-based embeddings. Additional comparisons with existing methods, including FuzDrop, DeePhase, catGRANULE2, PSPHunter, PhaSePred, SaPS, and Seq2Phase Scaffold, are also presented. **E)** PR-AUC versus ROC AUC for PS-Self vs. PS-Part classification on the independent test set. This analysis compares our downstream PS-Self and PS-Part models against other methods for classifying the two LLPS subtypes. **F)** ROC curves on the validation set for PS-Part vs. non-LLPS classification, comparing our ensemble PS-Part model with ESM2, SaProt, FuzDrop, DeePhase, catGRANULE2, PSPHunter, PhaSePred PdPS, and Seq2Phase Client.

A systematic evaluation and comparison of our phase separation protein classifiers demonstrated improved performance over existing methods. Our classifiers showed superior performance in classifying LLPS proteins compared to other methods. For LLPS vs. non-LLPS classification, our ensemble model achieved an area under the receiver operating characteristic curve (AUC) of 0.94 on the validation set, outperforming both of its individual components (ESM2: 0.93; SaProt: 0.91) and exceeding existing methods, including DeePhase (0.89), FuzDrop (0.85), PSPHunter (0.69), and catGRANULE2 (0.70) (**Fig. 2B**). On the external benchmarking test set, our ensemble model maintained higher performance with an AUC of 0.88 and an area under the precision-recall curve (PR-AUC) of 0.72, again surpassing existing methods (**Fig. 2C**). We next evaluated our classifiers’ ability to identify different LLPS protein subtypes. For PS-Self vs. non-LLPS classification, our ensemble PS-Self model achieved an AUC of 0.93 on the validation set, matching the performance of FuzDrop (0.93) and approaching the performance of DeePhase (0.95) and PhaSePred-SaPS (0.94) (**Fig. 2D**). The ESM2-based classifier (AUC of 0.93) outperformed the SaProt-based classifier (0.89), and the ensemble PS-Self matched the stronger of the two components for identifying PS-Self proteins (**Fig. 2D**). In contrast, existing methods showed limited ability to identify PS-Part proteins, likely because they were primarily trained on PS-Self datasets. For the PS-Part vs. non-LLPS classification task, our ensemble model achieved an AUC of 0.93 on the validation set, higher than DeePhase (0.82) and all other methods (AUC < 0.80) (**Fig. 2F**). Here, the SaProt-based classifier (0.93) slightly outperformed the ESM2-based classifier (0.92), and the ensemble model again matched the stronger of the two components (**Fig. 2F**). These findings highlight the importance of structural context for identifying PS-Part proteins and demonstrate a clear improvement over existing methods. Our ensemble classifiers can also effectively distinguish PS-Self proteins from PS-Part proteins. On the external test dataset (PS-Self vs. PS-Part), the ensemble model outperformed PhaSePred and Seq2Phase, two existing methods designed specifically for LLPS subtype classification, by over 3% in AUC and more than 5% in PR-AUC (**Fig. 2E**).

### Identifying critical regions promoting protein phase separation

Most data-driven machine learning methods for LLPS prediction often lack interpretability, offering little insight into which sequence features drive their predictions. Therefore, we evaluated whether our ensemble model could identify sequence regions that promote phase separation using a sliding window approach (**Fig. 3A**). Specifically, we divided protein sequences into overlapping windows of 21 residues to generate a positional LLPS propensity profile and defined “critical regions” as contiguous segments that exceeded a score threshold (**Fig. 3A**). We compared predicted critical regions with experimentally validated phase-separating regions from the PhaSePro database^43^ to evaluate the biological validity of these predictions. Fisher’s exact tests confirmed the significant enrichment of experimentally validated phase-separating regions within predicted segments for both PS-Self proteins (OR = 4.5) and PS-Part proteins (OR = 1.8; **Fig. 3B**). Among 49 PS-Self proteins, the ensemble model successfully recovered 85% of known phase-separating regions (50 of 59; **Fig. 3C**). For the more context-dependent PS-Part proteins, the predicted regions overlapped with 54% of known phase-separating regions (38 of 71; **Fig. 3D**). The comparatively stronger performance of the model in identifying critical regions of PS-Self proteins is consistent with their ability to phase-separate autonomously. In contrast, relatively modest performance among PS-Part proteins can be attributed to the requirement for partner interactions that lie outside any single-protein input. Nonetheless, our model’s ability to recover a significant number of PS-Part critical regions suggests it captures the intrinsic sequence and structural determinants these proteins contribute, independent of the partners that recruit them. Importantly, the predicted critical regions overlapped with many well-characterized phase-separation-driving elements across both protein classes (PS-Self and PS-Part) that mediate formation of biomolecular condensates (**Fig. 3E, 3G-H; Supplementary Fig. S2**). These included the low-complexity domain and RGG motifs of HNRNPA1^45,46^, the N-terminal and prion-like C-terminal domains of TDP-43^47,48^, the RNA recognition motifs of TIA1^49^, the RGG motif of FMRP^50^, and the prion-like low-complexity domains of EWS^51^. Our predictions also uncovered mechanistically distinct drivers, including the SH3 domains of GRB2^52^ and the disordered acidic-tract linker of NPM1^53^.

**Figure 3:**
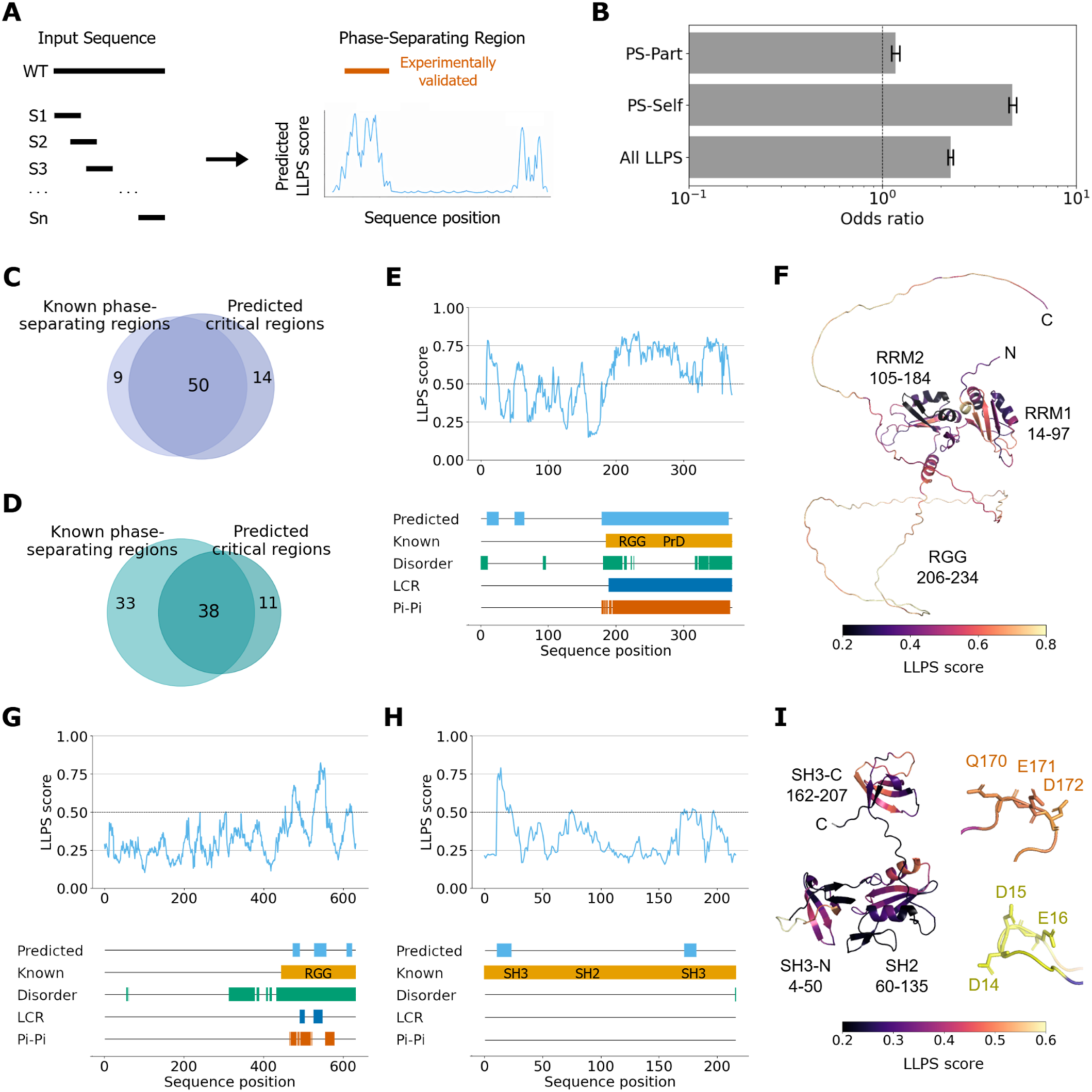
Identification and characterization of critical regions promoting phase separation. **A)** Schematic of the sliding-window method for generating residue-level LLPS propensity profiles of proteins. Consecutive sequence windows are scored by our ensemble model, and the resulting per-residue scores are binarized for comparison with known regions that promote phase separation. **B)** Odds ratio tests assessing enrichment of known phase-separating regions within predicted critical regions. Numbers of overlaps between known phase-separating regions and predicted critical regions in **C)** PS-Self and **D)** PS-Self proteins. **E)** Residue-level LLPS propensity profiles of HNRNPA1 (PS-Self) mapped alongside biochemical features. From top to bottom, the tracks show predicted critical regions, experimentally identified phase-separating regions, intrinsic disorder, low-complexity regions (LCRs), and predicted propensity for π-π interactions. **F)** AlphaFold2-predicted structure of HNRNPA1, colored according to residue-level LLPS propensity scores. **G)** Residue-level LLPS propensity profiles of FMRP (PS-Part). H) Residue-level LLPS propensity profiles of GRB2 (PS-Part). **I)** AlphaFold2-predicted structure of GRB2, colored according to residue-level LLPS propensity scores.

Beyond correctly identifying these well-characterized critical regions, our framework uncovered a small set of residues that drive our predictions, providing finer-grained, residue-level insight into how individual amino acids contribute to phase separation. For example, in the GRB2 protein, the negatively charged cluster D14/D15/E16 in the C-terminal SH3 domain and the Q170/E171/D172 cluster in the N-terminal SH3 domain were both identified as critical residues that promote phase separation (**Fig. 3I**), consistent with a previous study^54^. Mapping these predictions onto the corresponding AlphaFold-predicted structures further showed that the high-scoring residues localize to surface-exposed, disordered, or loop regions (**Fig. 3F & 3I**), consistent with their critical role in mediating multivalent intermolecular contacts. Taken together, these results demonstrate that our predicted critical regions closely align with experimentally identified phase-separating regions. Moreover, this suggests that our ensemble model captures biologically meaningful sequence determinants of phase separation and offers an interpretable, residue-level view of the underlying drivers.

### Proteome-wide enrichment & biochemical features analyses

To characterize the global pattern of phase-separating proteins, we applied our ensemble classifiers to the entire human proteome alongside established LLPS prediction methods, including FuzDrop^29^, PSPHunter^33^, DeePhase^35^, and catGRANULE2^34^. Our classifiers indicated that phase separation is likely widespread at the proteome level, with 7,514 proteins (37%) showing a high propensity to phase separate. Among these proteins, 2487 (12% of the proteome) were classified as PS-self and 5049 (25%) as PS-Part. Across all five methods, 2,331 proteins were consistently classified as LLPS, and 5,771 were consistently categorized as non-LLPS (**Supplementary Fig. S3A**). Our ensemble model uniquely predicted 162 proteins as LLPS, whereas other methods classified them as non-LLPS. Functional enrichment analysis of the consistently predicted LLPS proteins revealed strong enrichment for molecular interaction and cellular organization functions, with prominent localization to the nucleoplasm (**Supplementary Fig. S3B**). To further assess proteome-wide predictive behavior, we estimated lower-bound classification performance using proteins annotated in existing LLPS databases as positives and all remaining proteins as negatives. Under this conservative setting, our ensemble model maintained strong performance across the human proteome (AUC = 0.74), matching the sequence-based ESM2 model (AUC = 0.74) and outperforming SaProt (AUC = 0.72), PSPHunter (AUC = 0.68), FuzDrop (AUC = 0.56), and DeePhase (AUC = 0.55), while remaining below catGRANULE2 (AUC = 0.82) (**Supplementary Fig. S3C**).

To further characterize the LLPS proteins predicted by our models, we analyzed their subcellular localization patterns. We observed that proteins with high LLPS propensity scores, as quantified by our ensemble model, were strongly enriched in known biomolecular condensates, including stress granules, nuclear speckles, P-bodies, nucleoli, and nucleoplasm (**Fig. 4A**). These compartments are well known to host dynamic, membrane-less assemblies formed through phase separation. In contrast, proteins localized to membrane-bound organelles, such as mitochondria, lysosomes, the Golgi apparatus, peroxisomes, and lipid droplets, showed low LLPS propensity scores. This observation is consistent with membrane-bound organelles being structurally compartmentalized by lipid membranes and therefore relying less on dynamic, multivalent interactions. Beyond investigating localization patterns, we performed pathway enrichment analysis on the top-scoring 1,050 PS-Self and 1,664 PS-Part proteins to investigate their functional roles. We found that both PS-Self and PS-Part proteins were significantly enriched in the regulation of RNA metabolic processes (266 PS-Self genes, adjusted p-value 1e-58; 66 PS-Part genes, adjusted p = 1e-26) and cellular component organization (102 PS-Self genes, adjusted p = 1e-14; 530 PS-Part genes, adjusted p = 1e-46; **Fig. 4C**). Additionally, PS-Self proteins showed particularly strong enrichment in nuclear functions (**Fig. 4B & 4D**), consistent with their central role in nucleic acid-mediated condensation and scaffold formation. In contrast, PS-Part proteins exhibited broader enrichment patterns, spanning both nuclear functions (including chromatin and mRNA binding) and cytosolic functions (including cytoskeletal protein and adenyl ribonucleotide binding; **Fig. 4B & 4D**). Together, these findings show that our predictions recover well-established cellular localization and functional signatures of LLPS proteins in the human proteome.

**Figure 4.**
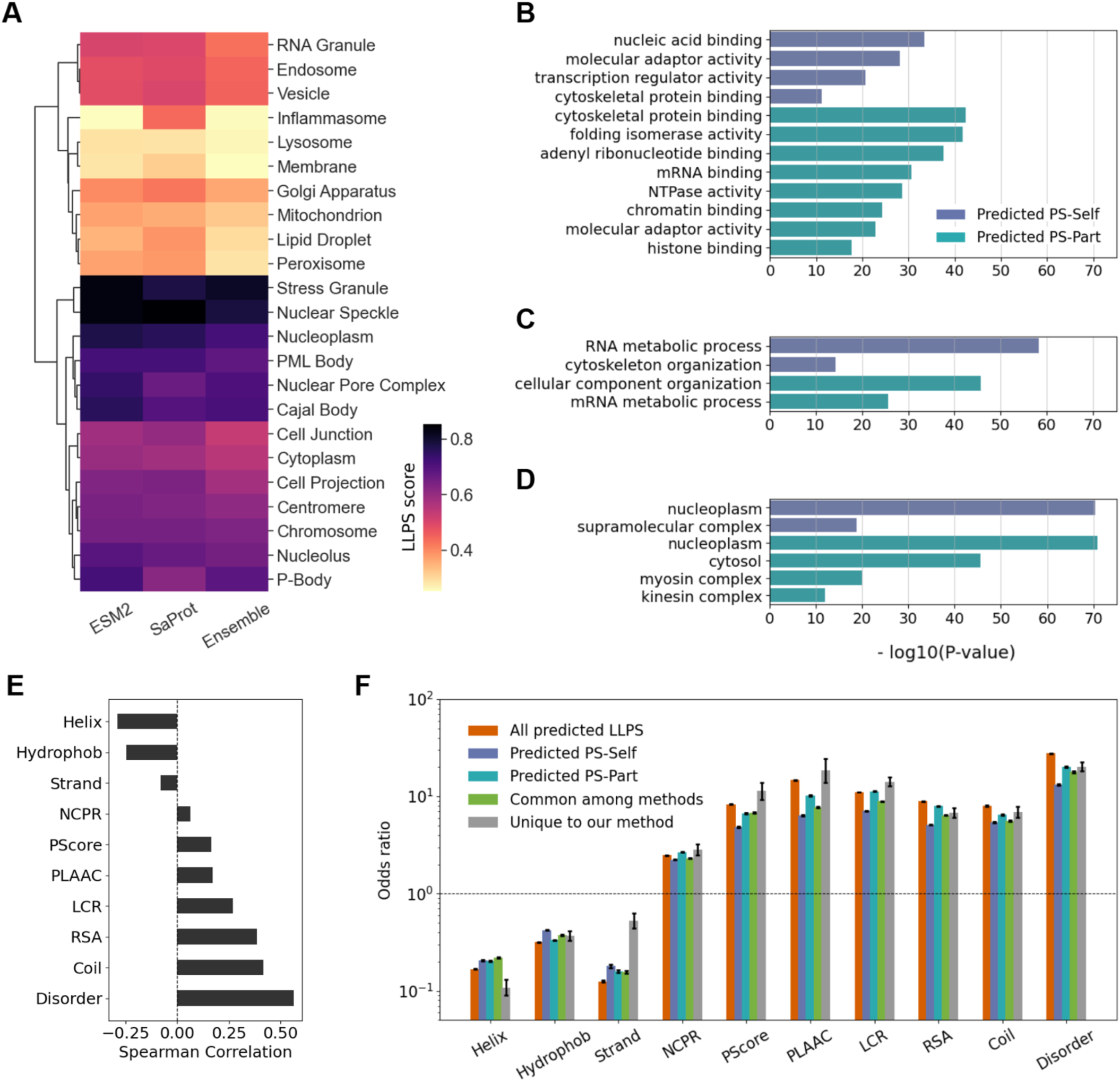
Proteome-wide enrichment analyses and biochemical features associated with predicted LLPS proteins. **A)** Subcellular localization analysis of the human proteome, showing LLPS propensity scores for proteins mapped to each cellular compartment. Black and yellow colors indicate higher and lower mean LLPS propensity scores across proteins within a particular subcellular location. Enrichment analyses of the top-scoring PS-Self and PS-Part proteins in the human proteome, showing enrichment of molecular functions (**B**), biological pathways (**C)**, and cellular components (**D)**. Purple and green colors correspond to enrichment scores for PS-Self and PS-Part proteins, respectively. **E)** Correlations between LLPS propensity scores and biochemical features computed across the proteome. **F)** Odds ratio tests evaluating enrichment of biochemical features in critical regions predicted by our ensemble model. Different colors in the bar plot correspond to distinct categories of LLPS proteins.

To assess the biological relevance of the predicted LLPS proteins, we correlated the predicted propensity scores with a range of known biochemical features that play critical roles in phase separation (**Fig. 4E**). We observed that intrinsic disorder exhibited a strong correlation with predicted LLPS scores (Spearman ρ = 0.56), consistent with the established role of disordered regions in promoting multivalent interactions that drive phase separation. Similarly, coil structure (ρ = 0.42) and relative solvent accessibility (RSA; ρ = 0.39) showed substantial positive correlations with predicted LLPS scores, reflecting the key role of flexible, solvent-exposed segments in facilitating dynamic interactions rather than forming stable folded structures. Interestingly, we observed moderate positive correlations between the predicted LLPS propensity score and other features, including low-complexity regions (LCR; ρ = 0.27), PScore (ρ = 0.17), and PLAAC (ρ = 0.17). In contrast, structural features characteristic of ordered protein cores were negatively correlated with LLPS propensity scores, including α-helix (ρ = -0.29) and hydrophobicity (ρ = -0.25).

Additionally, we analyzed residue-level enrichment patterns across biochemical features in predicted critical regions that significantly influence the model’s predictions (**Fig. 4F**). Consistent with protein-level correlations, the disorder feature showed the strongest enrichment (OR = 27.6) in the predicted critical regions of LLPS proteins. Predicted critical regions were also enriched for coil structure, RSA, low-complexity regions, PScore, and PLAAC (all OR > 1). In contrast, residues with high hydrophobicity and well-defined secondary-structure elements, including α- helices and β-strands, were significantly depleted (OR < 1) in these regions. Notably, LLPS proteins uniquely identified by our model showed residue-level biochemical profiles in critical regions similar to those identified by all methods, except for a modest increase in β-strand content. Overall, these observations indicate that the critical regions identified by our model recapitulate key biochemical features known to promote phase separation.

### Quantifying molecular impact of missense mutations on LLPS propensity

Coding mutations can alter the propensity of proteins to phase separate, disrupting associated biomolecular condensates and contributing to various diseases. Therefore, we systematically assessed the molecular impact of 85,263 missense mutations from the ClinVar and the denovo-db databases on the LLPS propensity of the underlying proteins. Our dataset included 28,006 benign mutations and 57,257 pathogenic mutations, spanning various disease phenotypes. We mapped these mutations to 1,539 predicted PS-Self proteins, 3,541 predicted PS-Part proteins, and 6,009 predicted non-LLPS proteins. The majority of missense mutations were sourced from ClinVar, with only 4.4% originating exclusively from the denovo-db and 0.2% shared between the two datasets (**Fig. 5A**). In our dataset, ClinVar contributed a larger proportion of benign mutations, whereas the Denovo-db entries were predominantly germline *de novo* mutations implicated in disease (**Fig. 5B**). To quantify the effects of these mutations on phase separation propensity, we applied our ensemble LLPS classifier to both wild-type and mutant sequences to compute the ΔLLPS score, defined as the change in predicted LLPS propensity upon mutation.

**Figure 5.**
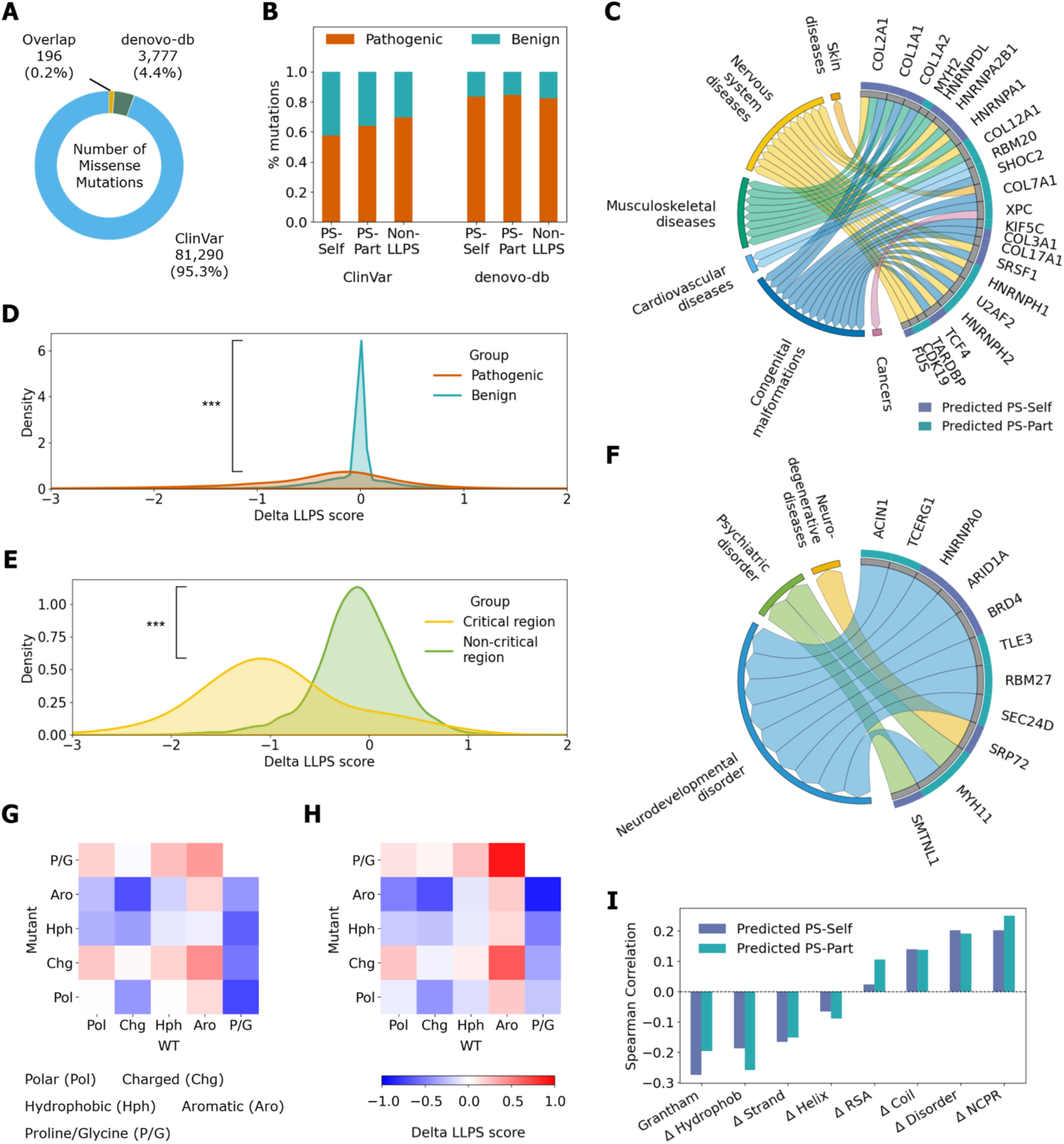
Disease-associated mutations from ClinVar and denovo-db that influence the LLPS propensity score. **A)** Numbers of missense mutations compiled from ClinVar and denovo-db. **B)** Percentages of pathogenic and benign mutations in our curated mutation dataset. **C)** Top-ranking disease-associated genes from ClinVar identified by combining high predicted LLPS propensity with enrichment of pathogenic mutations in predicted critical regions. **D)** Predicted changes in LLPS propensity (Delta LLPS) show a significantly broader distribution for pathogenic mutations than for benign mutations (two-sided Wilcoxon test, ***p < 0.001). **E)** Delta LLPS values are significantly lower for pathogenic mutations within predicted critical regions than for those outside these regions (two-sided Wilcoxon test, ***p < 0.001). **F)** Top-ranking disease-associated genes from denovo-db identified by combining high predicted LLPS propensity with enrichment of pathogenic mutations in predicted critical regions. Delta LLPS values for pathogenic mutations from ClinVar **(G)** and denovo-db **(H)** stratified by amino acid classes, including polar, charged, hydrophobic, aromatic, and Proline/Glycine. **I)** Correlations between mutation-induced changes in LLPS propensity (Delta LLPS) and corresponding changes in biochemical features.

As expected, we observed that benign mutations had ΔLLPS scores centered around zero, indicating minimal impact on phase-separation propensity. In contrast, pathogenic mutations exhibited a wider range of ΔLLPS scores, spanning both positive and negative values, which implies their increased potential to disrupt the phase-separation properties of affected proteins (Wilcoxon two-sided p < 0.001; **Fig. 5D**). These effects were even more evident when pathogenic mutations occurred within the critical regions predicted to promote phase separation. For instance, mutations within these regions showed a more pronounced reduction in LLPS propensity than those outside these regions (p < 0.001; **Fig. 5E**), indicating that mutations in these key sequence segments are more likely to disrupt the phase-separation behavior of the affected proteins. Together, these results demonstrate that our framework not only detects mutational impact but also highlights the sequence regions most likely to perturb phase-separation-associated function upon mutation.

In addition, we characterized residue-level physicochemical differences induced by mutations based on their effects on LLPS propensity. Previous studies have shown that both disorder-promoting residues (glycine and proline) and aromatic residues play key roles in mediating the weak, multivalent interactions that drive phase separation^33,47^. These roles are often described within the framework of stickers and spacers, in which aromatic and charged residues act as interaction stickers embedded within flexible, disordered spacer regions. Consistent with this framework, we found that missense mutations that perturb disorder-promoting residues showed a greater decrease in ΔLLPS scores than other mutations, indicating a stronger loss in phase-separation propensity (**Fig. 5G & 5H**). Similarly, missense mutations that convert wild-type charged amino acid residues to aromatic residues strongly reduced the propensity for phase separation (**Fig. 5G & 5H**), likely by disrupting charge clusters and altering cation-pi or pi-pi interactions. Furthermore, we compared ΔLLPS scores with mutation-induced changes in biochemical features between wild-type and mutant proteins (**Fig. 5I**). Changes in net charge per residue (NCPR) and disorder showed weak positive correlations with ΔLLPS scores (Spearman ρ = 0.20 and 0.23, respectively), indicating that reduced LLPS propensity is associated with decreases in these features. Conversely, the Grantham score is negatively correlated with ΔLLPS scores (ρ = -0.24), indicating that mutants with greater evolutionary distance from their wild types tend to disrupt phase separation. These results are consistent with prior studies indicating the critical roles of IDRs and electrostatic interactions in maintaining the phase-separation behavior of proteins^46,55,56^. Interestingly, changes in RSA and hydrophobicity showed stronger correlations with ΔLLPS in PS-Part proteins than in PS-Self proteins, reflecting underlying differences in the biochemical composition and sequence determinants of phase separation between the two LLPS subtypes (**Fig. 5I**).

We performed a systematic analysis of disease-associated missense mutations across 11,109 human genes and ranked them by their association with aberrant protein phase separation. Among proteins encoded by these genes, our ensemble model annotated 1,539 as PS-Self, 3,541 as PS-Part, and 6,009 as non-LLPS, providing the basis for subsequent disease-association analyses. By ranking changes in protein phase-separation propensity, critical region coverage, and pathogenic missense mutation burden (from ClinVar and denovo-db), we sought to identify genes in which aberrant phase separation is most likely to contribute to disease. This comprehensive analysis identified many well-established LLPS proteins for which aberrant phase separation has been linked to various diseases. Among the top-ranking proteins based on ClinVar data, many were RNA-binding proteins, including FUS, TDP-43, RBM20, HNRNPA1, and HNRNPD, all of which are associated with various pathological conditions (**Fig. 5C**). The most prevalent disease classes associated with these proteins included congenital malformations, nervous system disorders, and musculoskeletal diseases. At the level of specific disorders, neurodegenerative diseases, including Amyotrophic Lateral Sclerosis and Frontotemporal dementia, were the most prominent, followed by neurodevelopmental disorders, inherited disorders, muscular dystrophy, and muscular dysplasia. The top-ranking proteins based on the denovo-db data included BRD4 and HNRNPA0, and the most-represented disease class was neurodevelopmental disorder (**Fig. 5F**). The top disorders driven by de novo mutations affecting LLPS proteins included Autism, Tourette Syndrome, and Amyotrophic Lateral Sclerosis.

### Examples of disease-associated pathogenic mutations predicted to disrupt phase separation

After investigating global patterns of how mutations modulate phase-separation propensity, we next characterized individual LLPS proteins harboring known pathogenic mutations implicated in various diseases. Across well-established LLPS proteins, pathogenic mutations frequently clustered within critical regions predicted by our framework to drive phase separation. For example, pathogenic mutations frequently occur in the predicted critical region of the C-terminal prion-like domain in the TDP-43 protein (**Fig. 6B**). Similarly, many Rett syndrome-associated mutations mapped to the methyl-CpG-binding domain (MBD), which we identified as a critical region for the MECP2 protein (**Supplementary Fig. S4B**). Our framework also highlighted the prion-like domain of HNRNPA1 as a critical region, with pathogenic mutations associated with amyotrophic lateral sclerosis and multisystem proteinopathy affecting this region (**Supplementary Fig. S4D**). These observations suggest that disease-associated mutations in LLPS proteins preferentially occur in regions that contribute directly to condensate formation and regulation.

**Figure 6.**
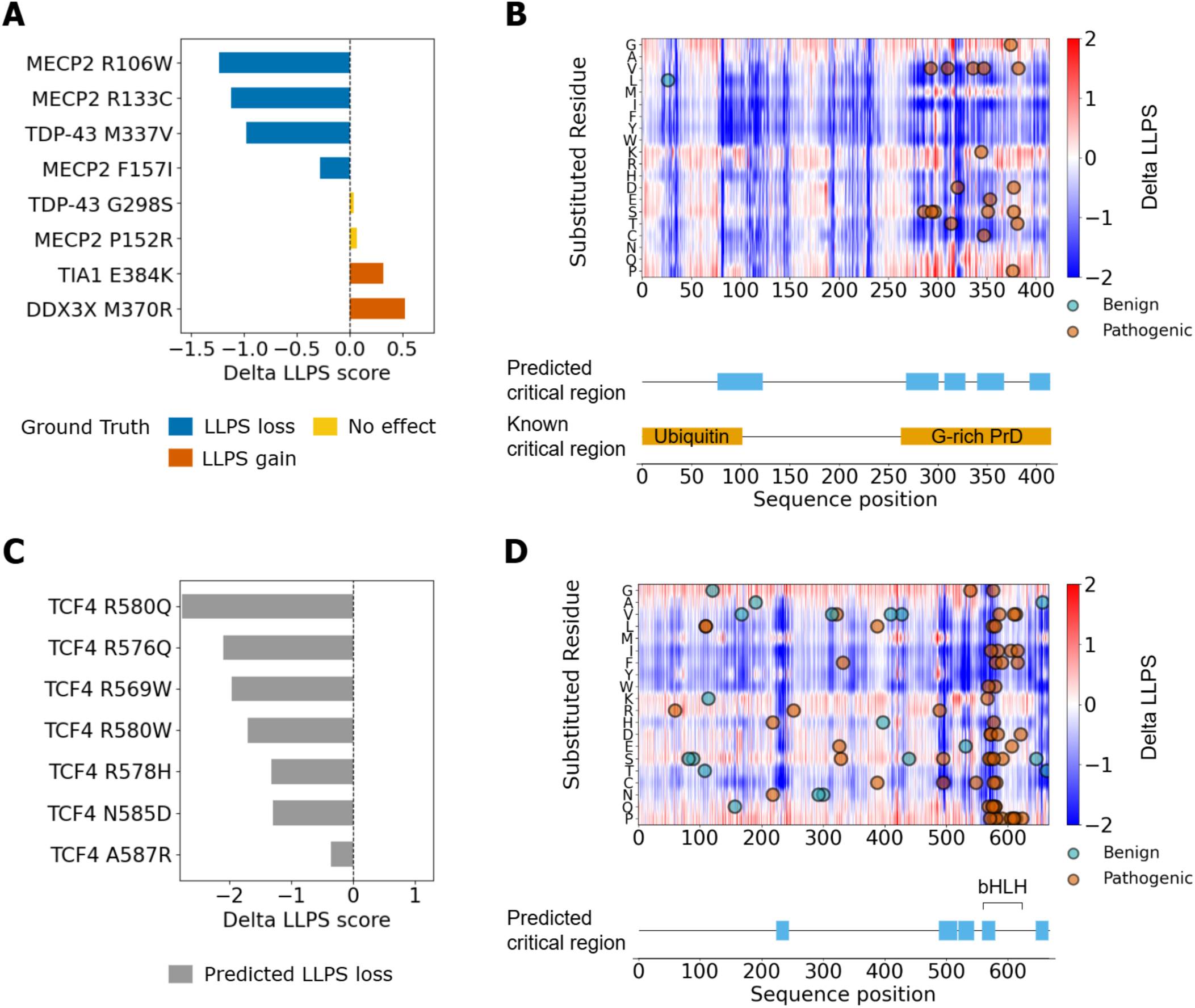
Example of mutational impact on phase-separation propensity for well-known and novel predicted LLPS proteins. **A)** Predicted effects on LLPS propensity (Delta LLPS) for a set of well-characterized pathogenic mutations in experimentally validated LLPS proteins. **B)** Saturation mutagenesis heatmap for the TDP-43 protein, showing the predicted effects of all single mutants on LLPS propensity. The blue and red colors on the heatmap indicate positive and negative Delta LLPS values, respectively. Overlaid points represent missense mutations (cyan for benign and brown for pathogenic) from the ClinVar and denovo-db databases. The tracks above the heatmap indicate predicted critical regions (blue) and known phase-separating regions (orange). **C)** Predicted effects of Pitt-Hopkins mutations on LLPS propensity for the candidate LLPS protein TCF4. **D)** Saturation mutagenesis heatmap for TCF4, showing the predicted effects of all single mutants on LLPS propensity.

Our framework accurately captured pathogenic mutations that reduce or preserve LLPS (**Fig. 6A**). For instance, the ALS-associated mutation M337V in the TDP-43 protein resulted in a strong decrease in phase-separation propensity (ΔLLPS = -0.98, z-score = -1.36), consistent with experimental studies that these mutations impair LLPS and enhance pathological aggregation^16^. In contrast, the ALS-associated mutation G298S in this protein showed a minimal predicted effect on LLPS propensity (ΔLLPS = 0.04, z-score = 0.35), consistent with previous experimental studies^54^. Similarly, several Rett syndrome-associated mutations in the MBD region of the MECP2 protein, including R106W, R133C, and F157I, were predicted to substantially reduce phase-separation propensity (ΔLLPS = -1.24, -1.12, and -0.28; z-scores = -1.78, -1.59, and -0.20, respectively). These predictions are consistent with studies showing that mutations in the MBD region of MECP2 impair chromatin condensation and weaken LLPS behavior by disrupting interactions with methylated DNA^55^. Similarly, our framework correctly recovered experimental findings^55^ that the Rett syndrome-associated mutation P152R has minimal effect on LLPS propensity (ΔLLPS = 0.07, z-score = 0.39).

Beyond identifying mutations that reduce phase-separation propensity, our method also identified pathogenic mutations that increase propensity scores. For the stress granule protein TIA1, although the full-length protein is annotated in PhaSePro as critical for phase separation, our framework highlighted the IDRs and RNA recognition motif domains as critical regions (**Supplementary Fig. S4A**). In particular, the Welander distal myopathy-associated mutation E384K was predicted to increase LLPS propensity (ΔLLPS = 0.32, z-score = 0.80), consistent with studies showing enhanced phase separation and impaired stress granule disassembly caused by this mutation^49^. Similarly, the DDX3X mutation M370R, implicated in multiple cancer types including medulloblastoma and chronic lymphocytic leukemia, showed increased LLPS propensity (ΔLLPS = 0.53, z-score = 1.15; **Supplementary Fig. S4C**). Previous studies suggest that this mutation promotes stress granule assembly and stabilizes condensate formation^57^.

Furthermore, our framework identified high-ranking novel or emerging candidate LLPS proteins that lack experimental validation and may contribute to disease through altered phase-separation behavior. For instance, our model assigned a high LLPS propensity score (0.97) to TCF4, a transcription factor associated with neurodevelopmental disorders and schizophrenia (**Fig. 6D**). We observed that a predicted critical region (residues 487-566) within TCF4 was highly enriched for disorder and low-complexity features. In particular, the Pitt-Hopkins disease-associated mutations R569W, N585D, and A587R in this critical region yielded negative ΔLLPS scores of -1.96, -1.29, and -0.36, respectively (**Fig. 6C**), suggesting that disruption of phase separation may contribute to this disease. Together, these examples highlight that critical regions for phase separation identified by our framework not only match known phase-separating domains but also overlap with sites of disease-associated mutations. Additionally, our framework’s ability to quantify residue-level ΔLLPS scores enables mechanistic interpretation of how single-point mutations modulate phase separation behavior, providing insights into their pathogenic impacts.

## Discussion

Comprehensive interpretation of how coding variants affecting IDRs contribute to disease remains challenging, as most variant-effect prediction tools were developed for structured protein regions and therefore provide limited mechanistic insight into mutations in disordered sequences. Evaluating perturbations in IDR-mediated phase-separation propensity offers an attractive approach to addressing this gap. However, achieving this at scale requires accurate, interpretable LLPS predictors that can resolve distinct modes of phase separation. Therefore, we developed an ensemble machine-learning framework that uses sequence and structural embeddings from protein-language models to predict the propensity for phase separation from protein sequence and structure. Our framework operates in two steps, where the first classifier distinguishes LLPS from non-LLPS proteins, and downstream classifiers further classify LLPS subtypes (PS-Self and PS-Part). Overall, our model outperformed existing methods across all three classification tasks, with the largest gains observed for the PS-Part class in independent benchmark datasets. Furthermore, the interpretable approach of our model enabled us to identify well-known and novel pathogenic missense mutations from ClinVar and the denovo-db that are likely to play critical roles in disease etiology by promoting aberrant phase separation.

Establishing a mechanistic link between pathogenic mutations and human disease through aberrant phase separation requires insight into the physicochemical properties of residues and regions that drive such predictions. Therefore, we emphasized model interpretability in our framework to pinpoint the regions that contribute most to the model’s output, labeling them as critical regions in individual proteins that align with experimentally validated phase-separating regions. Our approach differs from prior data-driven LLPS predictors^30,35,41^, which often lack such interpretability. While some recent tools have focused on improving interpretability, significant limitations remain. For instance, PSPHunter^33^ identifies key residues driving phase separation by masking each protein sequence segment, but the sensitivity of the resulting profile is affected by protein length and requires manual parameter adjustment. Similarly, the catGranule2^34^ predictor highlights key residues by training a separate model on shorter segments. However, its residue-level outputs are not directly tied to the model that produces the propensity score. In contrast, our framework provides a direct interpretation of the predicted propensity score by identifying the underlying regions using the same model, eliminating the need for post-hoc parameter tuning or auxiliary models. Furthermore, our systematic interpretability analyses enabled us to identify various biophysical characteristics of critical residues and sub-regions consistent with established LLPS biophysics. For example, we found intrinsic disorder, π-π stacking interactions, and nucleic acid binding to be positive contributors to the propensity for phase separation. Moreover, PS-Self proteins show higher enrichment in IDRs and RNA-binding granules, whereas PS-Part proteins exhibit more structured regions, higher dependence on solvent accessibility, and enrichment in modular binding domains, signaling condensates, and chromatin/transcription hubs. Our approach of integrating predicted structural context with sequence information further enriched these predictions, leading to improved predictive performance of our ensemble model. This improved performance can be attributed to our ensemble model’s ability to better capture transient local structure and ensemble properties that sequence-only representations do not fully resolve.

Beyond interpretability, our ensemble method outperformed other LLPS detection methods likely due to the diversity of the underlying training data and an optimized model architecture. In particular, our model showed a significant gain in predictive performance over other methods when classifying PS-Part proteins versus PS-Self proteins. The low performance of existing methods in accurately classifying PS-Part proteins may be attributed to inherent bias in the training datasets, which are heavily focused on PS-Self proteins, limiting these models’ ability to learn heterotypic interaction signals critical for identifying PS-Part proteins. To address this limitation, we used PS-Self and PS-Part proteins from various LLPS databases, as well as a proteome-wide experimental study, to achieve more balanced coverage of both modes of phase separation. Furthermore, existing supervised machine-learning-based LLPS predictors primarily use fully structured proteins from the PDB as negative labels and often overlook disordered non-LLPS proteins, thereby conflating “non-phase-separating” with “structured” and inflating apparent performance on disordered test proteins. Therefore, we included both fully structured and disordered proteins in our negative set by selecting proteins that are not known to interact with LLPS proteins. This strategy reduced the risk of training the model to simply distinguish between structured and disordered regions and produced a more rigorous benchmark for what the model learns about phase separation. Additionally, our approach of integrating predicted structural embeddings with sequence representations provided additional information about transient local structure and ensemble properties, complementing the sequence signal and contributing to the performance gains observed for both PS-Self and PS-Part classification.

To evaluate the efficacy of our model for predicting the impact of mutations affecting IDRs, we applied our framework to systematically analyze phase-separation perturbations induced by disease-associated variants in ClinVar and the denovo-db. Across 81,486 missense variants in ClinVar, we observed 2,025 pathogenic or likely pathogenic mutations mapping to critical regions of predicted LLPS proteins and showing meaningful changes in phase-separation propensity. Our analyses identified both established and novel LLPS proteins for which aberrant phase separation offers a plausible mechanistic link to disease. For instance, a previous study has shown that the ALS-associated M337V mutation in TDP-43 falls within the conserved α-helical segment of the prion-like C-terminal LCD that mediates helix–helix assembly, thereby weakening these contacts and biasing the system toward irreversible aggregates^47^. Consistent with this finding, our method predicted a decrease in phase-separation propensity induced by this mutation. In contrast, we observed an increase in predicted LLPS propensity induced by the medulloblastoma-associated mutation in the DDX3X protein. This observation aligns with a prior study indicating that M370R introduces a positively charged arginine residue that strengthens electrostatic and cation–π contacts with adjacent LCD and RG/RGG regions, resulting in stress-granule hyper-assembly^57^. Similarly, the Rett-syndrome mutation R133C in MECP2, which removes a guanidinium group that hydrogen-bonds with methylated CpG DNA and thereby couples the loss of DNA binding to the loss of the methylation-dependent multivalency that drives MECP2 condensation^58^, is in line with our predicted reduction in phase-separation propensity. Beyond recapitulating known cases, our framework identifies TCF4 as a novel/emerging LLPS candidate, with Pitt–Hopkins pathogenic variants mapping to predicted critical regions of this DNA-dependent phase-separating transcription factor^59^. Overall, our approach can detect both putative loss-of-LLPS (TDP-43, MECP2) and gain-of-LLPS (DDX3X) phenotypes, suggesting that our framework can capture bidirectional perturbations in condensate behavior. More broadly, these examples implicate aberrant condensate behavior as a previously unrecognized disease mechanism in these and other genes. We note that these predictions complement rather than replace existing VEPs that primarily focus on structural stability, including AlphaMissense^60^, EVE^61^, and ESM1B-based VEPs^62^. Although these structure-stability-focused VEPs can still assign pathogenicity scores to IDR variants, they lack mechanistic insight into how these mutations might contribute to pathogenicity within IDRs. Our framework addresses this mechanistic gap by attributing variant effects to perturbations in the propensity for phase separation.

Despite improved predictive performance, greater interpretability, and demonstrated utility in linking potential aberrant phase separation to disease mechanisms, we note several limitations in our approach that should be considered when interpreting our framework’s outputs. In particular, experimentally validating the predicted impact of mutations on LLPS propensity is a significant challenge, as such validation primarily depends on the environmental and cellular conditions that govern the protein of interest’s LLPS characteristics. Additionally, missense mutation-induced changes in LLPS propensity alone cannot establish causality, since IDR variants may simultaneously affect phase separation, short linear motifs, and post-translational modification sites. Therefore, disentangling these distinct contributions requires targeted experimental follow-up. Notably, despite accurately predicting LLPS propensity, our method does not characterize the material properties of condensates, which can adopt distinct states (liquid-like, gel-like, or solid-like) or undergo protein aggregation. Insight into pathological transitions between these states (e.g., liquid-to-solid) and aggregation processes is critical for elucidating mechanisms underlying several neurodegenerative diseases. Furthermore, environmental factors, including pH, ionic strength, temperature, and protein concentration, can significantly influence the conditions under which phase separation occurs. However, our model assumes physiological conditions and does not predict LLPS propensity under specific environmental perturbations. This is a critical gap that needs to be addressed, as experimental studies have shown that modest changes in buffer conditions can dramatically alter phase-separation behavior, and that mutations may exert context-dependent effects not captured by a static prediction. Similarly, although our training set improves upon prior work in PS-Part and negative-set coverage, available LLPS databases still overrepresent certain protein families, particularly RNA-binding proteins and transcription factors, which might introduce bias in predictive performance across distinct protein families. Finally, our framework predicts the propensity of proteins to phase separate but not which specific bimolecular condensate they join, as many LLPS proteins participate in multiple condensates with distinct partners and functional roles.

There are several potential extensions to our current work to address the limitations outlined above. For instance, incorporating features that capture liquid-to-solid transitions will likely enable more direct modeling of pathological condensate behavior, which will be critical for investigating the role of aberrant condensates in neurodegenerative disease. Similarly, incorporating post-translational modifications that dynamically regulate condensate assembly and dissolution within our current framework will be important. Including PTMs will likely add a layer of condition-dependent prediction that is currently absent from most LLPS tools. Furthermore, beyond predicting LLPS propensity, assessing condensate composition and partner-specific recruitment will improve the accuracy of our mutational impact analyses and provide additional biophysical insights into the role of aberrant LLPS in various diseases. We also note that as experimental LLPS datasets expand through high-throughput approaches, retraining on broader, more balanced data will help address current biases stemming from protein-family coverage and improve our model’s generalization across the human proteome. Despite these limitations and the need to address them, our work overall demonstrated that phase separation offers a viable and interpretable framework for analyzing protein-coding variants in IDRs. We anticipate that as condensate biology continues to mature, mechanism-aware tools such as ours will likely bridge the current gap between computational variant prediction in IDRs and the identification of condensate-modulating therapeutic targets across various diseases^8–12^.

## Materials & Methods

### Curating LLPS and non-LLPS protein datasets

For model training and validation, we assembled a positive set of LLPS proteins and a negative set of proteins unlikely to undergo phase separation under physiological conditions. We compiled the positive set from a proteome-wide study and several public databases. For instance, we used data from a recent experimental study that identified 1,518 endogenous condensate proteins in a cell’s proteome by sorting proteins based on their oligomeric states^42^. These proteins were experimentally validated to participate in condensates but lack annotations distinguishing self-assembled phase-separating (PS-Self) proteins from partner-dependent phase-separating (PS-Part) proteins. In addition, we selected 150 PS-Self proteins annotated as “scaffold” and 8,161 PS-Part proteins annotated as “client” from the DrLLPS database^40^ (v1.0). Similarly, we obtained 347 PS-Self proteins, which are single-component phase-separating proteins, and 324 PS-Part proteins in multi-component systems from the LLPSDB database^39^ (v2.0). We obtained this subset by removing entries with post-translational modifications or single-site mutations. Additionally, we required phase separation to be observed at solute concentrations below 100 μM, indicating a strong propensity for phase separation. Finally, we selected 57 PS-Self proteins by excluding entries labeled as “partner-dependent”, “RNA-dependent”, or “PTM-dependent”, and 51 PS-Part proteins labeled as “partner-dependent” from the PhaSePro database^43^ (v1.1.0). After merging datasets from all these sources, we discarded sequences shorter than 50 amino acids and removed redundant sequences. Our final positive set included 4,857 LLPS proteins, comprising 748 experimentally confirmed LLPS proteins, 352 PS-Self proteins, and 3,757 PS-Part proteins. To construct the negative set of proteins unlikely to undergo phase separation, we removed all proteins in the positive set from the Protein Data Bank^63^ (PDB). We also removed their first interactors, which are potential phase-separation participants, using protein-protein interaction data from the BioGRID database^64^ (v3.5.175). From the remaining pool, we randomly selected 4,857 non-LLPS proteins to form our negative set.

### Construction of Training, Validation, and Test datasets

We constructed separate datasets for different classification tasks. The training set for the LLPS classifier (LLPS vs. non-LLPS) included a randomly sampled subset of 236 PS-Self proteins and 3,641 PS-Part proteins from the positive set, along with all 1,518 experimentally identified LLPS proteins^42^, 748 of which do not overlap with other proteins in the positive set (**Supplementary Table 1 & Supplementary Data S1-S3**). To balance the classes, we also sampled 4,625 non-LLPS proteins from the negative set. For training the PS-Self and PS-Part classifiers, we excluded the 748 experimentally identified LLPS proteins that lack explicit PS-Self/PS-Part annotations and trained only on the subset of proteins with curated PS-Self/PS-Part labels. For hyperparameter tuning and model evaluation, we constructed a validation set comprising the remaining 116 PS-Self proteins, 116 PS-Part proteins, and 232 non-LLPS proteins, none of which were used during training. The test set is an external benchmark dataset^44^ compiled from various databases, including CD-CODE^65^, DrLLPS^40^, LLPSDB^39^, PhaSepDB^38^, PhaSePro^43^, DisProt^66^, and PDB^63^. We removed all proteins present in the training set from the test set before evaluating our models’ performance. We used the performance metrics reported in the benchmark study^44^ to compare our approach with other methods.

### Ensemble model architecture for predicting LLPS proteins

We first trained two distinct models using different embeddings to predict LLPS proteins. The sequence-only model incorporated residue-level embeddings from the protein-sequence-based language model ESM2, whereas the structure-based model utilized embeddings from the protein-structure-based language model SaProt. We then trained an ensemble model that combines complementary information from these two embeddings. For sequence-based representations, we embedded each protein sequence using the ESM2 model (esm2_t33_650M_UR50D), which was pretrained on large-scale protein sequence databases and encodes information about protein composition, evolutionary constraints, and functional motifs^67^. In parallel, we incorporated structural context using the SaProt model (SaProt_650M_AF2), which augments sequence-based representations with local structural context^68^. To generate the structure-aware inputs required by SaProt, we processed all protein sequences using Foldseek^69^ (release 10), which encodes local backbone geometry and residue environments into discrete structure-aware tokens. SaProt then converts these tokens into residue-level structure-aware embeddings. For both the ESM2- and SaProt-based models, the embeddings were passed through a multi-head self-attention layer followed by a linear projection that outputs a single propensity score for each protein (**Supplementary Fig. S1A-B**).

Our ensemble model architecture captures both amino acid sequence patterns and 3D structural context by integrating two pretrained protein language models (**Fig. 1A**). Because both ESM2 and SaProt produce residue-level embeddings with a dimensionality of 1280, their embeddings can be directly fused after the self-attention layers. Specifically, we applied a multi-head cross-attention layer, in which the SaProt embeddings served as the queries and the ESM2 embeddings served as the key-value pairs. By conditioning structure-aware embeddings on rich sequence-level information, this fusion layer implicitly learns how structural context modulates sequence-based features. A final linear layer projects the resulting fused representation into a propensity score for each protein (**Supplementary Fig. S1C**). Unless otherwise stated, we report all downstream analyses and results using the ensemble model.

Notably, we generated structure-aware embeddings for the entire protein sequence, including both structured and IDR components. IDRs are an essential feature of many phase-separating proteins and often mediate multivalent, weak interactions that drive condensate formation. Since predicted structures are inherently less reliable in disordered regions, we expect the model to rely more heavily on sequence-based embeddings in these regions, while exploiting structural context primarily where stable secondary and tertiary structures are present (**Supplementary Table 2**). During training, we updated only the attention layers and the final linear layer, keeping all other model parameters frozen. This approach allows the models to learn with minimal additional training cost and reduced overfitting, particularly given the limited availability of experimentally validated LLPS data. We used the binary cross-entropy loss function, four attention heads per attention layer, a dropout rate of 0.6, the Adam optimizer (lr=1e-4), and a batch size of 16 to train all models. Training was stopped if the loss did not decrease for 5 epochs. We implemented this framework in PyTorch 2.9.1+cu126 and trained our models on GPUs (NVIDIA H100).

### Hierarchical classification workflow to detect PS-Self and PS-Part proteins

Our hierarchical classification workflow uses models with the same basic architecture but were trained separately on different datasets for each task. The LLPS classifier (LLPS vs. Non-LLPS) outputs a propensity score indicating the likelihood that a protein will undergo phase separation. Proteins with a score above 0.5 are labeled as LLPS and proceed to the PS-Self and PS-Part classifiers, while those below are considered non-LLPS (**Fig. 1B**). We chose the 0.5 threshold to maximize the F1 score on the validation set (**Supplementary Table 3**). The PS-Self and PS-Part classifiers further categorize LLPS proteins as either self-assembling or partner-dependent. Since some proteins can act as both PS-Self and PS-Part in different contexts, a single binary classifier is insufficient. Therefore, we trained two separate classifiers: one to identify PS-Self proteins and the other to identify PS-Part proteins. For the PS-Self classifier, all proteins annotated as self-assembling were labeled as positives, and all others were labeled as negatives. The output of this model is a PS-Self score indicating the probability that an LLPS protein undergoes self-assembly-based phase separation. Conversely, for the PS-Part classifier, all proteins annotated as partner-dependent were labeled as positives, and all others were labeled as negatives. The output from this model is again a PS-Part score, representing the probability that an LLPS protein undergoes partner-dependent phase separation. This approach allows the models to focus on specific subtypes and capture the overlapping roles of LLPS proteins.

### Using model interpretability to identify critical regions

We used a sliding window method to identify sequence segments that promote phase separation (**Fig. 2A**). A fixed-length window of 21 residues moved sequentially along each protein sequence and its corresponding structure-aware tokens. Our ensemble classifier (LLPS vs. non-LLPS) processed each window independently and assigned a propensity score to the central residue. For residues near the N- or C-termini, we truncated the window to accommodate sequence boundaries. Because the ensemble classifier was trained on full-length protein sequences, direct application to short sequence windows yields scores on a different scale. To account for this difference, we calibrated the per-residue scores using 22 LLPS proteins from the training set, with experimentally annotated phase-separating regions from the PhaSePro database^43^ (**Supplementary Method**). Residues with calibrated scores greater than 0.5 were defined as belonging to predicted critical regions. To obtain contiguous, non-trivial segments, we post-processed the predictions by filling gaps shorter than 11 residues between adjacent critical regions and removing isolated critical regions shorter than 11 residues. This sliding window method provides interpretability at the residue level by highlighting specific sequence segments most likely responsible for promoting phase separation. For downstream evaluation, we compared predicted critical regions with experimentally validated phase-separating regions from the PhaSePro database^43^. We applied the sliding window method to the remaining 49 PS-Self proteins and 42 PS-Part proteins annotated in PhaSePro. We then performed Fisher’s exact tests to assess enrichment of predicted critical regions within experimentally validated phase-separating regions. An odds ratio (OR) greater than 1 indicates enrichment of predicted critical regions in known phase-separating regions, whereas an OR less than 1 indicates depletion.

### Characterizing biochemical features of predicted critical regions

We computed a range of established biochemical features associated with phase separation at both the protein and residue levels to investigate the relationship between the predicted LLPS propensity and the underlying biochemical properties of proteins. For example, we quantified the intrinsic disorder score from the protein sequence using AIUPred^70^ (v1.2.2). Similarly, we calculated local sequence properties, including NCPR, fraction of charged residues (FCR), and hydrophobicity, using localCIDER^71^ (v0.1.20). We used the SEG^72^ method to identify low-complexity regions (LCRs) with a window size of 12 residues, K1 = 2.2 (threshold for initiating an LCR), and K2 = 2.5 (threshold for extending an LCR). Additional sequence determinants relevant to LLPS were captured using PScore^32^ (v2), which measures pi-pi interactions (contact distance between planar surfaces), and the PLAAC method^31^, which indicates the presence of prion-like domains. We incorporated structural features by predicting relative solvent accessibility (RSA) and secondary-structure elements, such as helix, strand, and coil, using NetSurfP^73^ (v3.0).

We compared averaged per-residue features across sequences with the predicted propensity scores to evaluate global correlations. We also conducted Fisher’s exact tests to assess whether the per-residue features are enriched in the predicted critical regions. An OR above 1 indicates enrichment of a feature in those regions, while an OR below 1 indicates depletion. These analyses not only assess the biological relevance of the predicted propensity scores but also identify the sequence and structural features that most strongly contribute to phase separation behavior.

### Cellular localization & pathway enrichment analyses of putative LLPS proteins

To investigate whether the propensity for phase separation varies across cellular compartments, we applied the ensemble classifier (LLPS vs. non-LLPS) to a proteome-wide dataset of 15,283 proteins (**Supplementary Data S4)** with annotated subcellular localizations^29^. This dataset included proteins associated with both membrane-bound organelles (e.g., mitochondrion, Golgi apparatus, endosome) and membrane-less organelles (e.g., nucleolus, stress granule, Cajal body). The ensemble classifier predicted a propensity score for each protein, which was then averaged across proteins within each cellular compartment to generate a compartment-level score. The overall goal of this analysis was to assess whether proteins localized to membrane-less organelles, often associated with phase-separated biomolecular condensates, exhibit higher predicted LLPS propensity than those in membrane-bound compartments. We performed pathway enrichment analyses of the top PS-Self and PS-Part proteins in the human proteome. First, we selected top-scoring PS-Self proteins with a predicted propensity score above 0.9 and a PS-Self score above 0.5. Similarly, we selected top-scoring PS-Part proteins with a predicted propensity score above 0.9 and a PS-Part score above 0.5. Next, we provided each resulting gene set independently as input to g:Profiler^74^ to identify enriched biological pathways using the default enrichment parameters. Finally, we retained only the most significant pathways with adjusted p-values below 1e-10.

### Genomic Variant Dataset Analyses

To investigate disease-associated mutations that may alter the propensity for protein phase separation, we assembled a high-confidence set of human genomic variants from the ClinVar^75^ (version 2025-10) and the denovo-db^76^ (v1.6.1) databases (**Supplementary Data S5)**. By comparing predicted propensity scores of wild-type and mutant protein sequences, we identified mutations that may disrupt phase separation. For the ClinVar data, we retained only entries mapped to the GRCh38 human reference genome and restricted the analysis to missense mutations, which directly alter amino-acid identity and can affect phase-separating regions. To ensure reliability, we included variants with one of the following review statuses: “criteria provided, single submitter,” “criteria provided, multiple submitters, no conflicts,” “reviewed by expert panel,” or “practice guideline.” We further filtered variants by clinical significance, keeping only those labeled “Pathogenic,” “Likely pathogenic,” or “Benign,” to balance the numbers of pathogenic and benign mutations. For the denovo-db data, all variants are mapped to the GRCh38 human reference genome. We included variants from both the Simons Simplex Collection (SSC) and non-SSC subsets and selected only validated missense mutations. We considered both disease-associated and “control” mutations to assess phase separation across pathogenic and benign de novo mutations.

### Amino Acid Composition Analyses and Mutation Effect Predictions

We categorized amino acids by their physicochemical properties and investigated their effects on phase separation. Specifically, residues were grouped into polar (S, T, C, N, Q, H), charged (D, E, K, R), hydrophobic (A, V, L, I, M), and aromatic (F, W, Y) categories. In addition, proline (P) and glycine (G) were treated as a separate category of disorder-promoting amino acids, given their established roles in intrinsic disorder and in flexible regions of proteins^16,21,77^. Previous studies have highlighted the enrichment of P and G in disordered regions and their involvement in pathogenic mutations, suggesting that these residues can modulate conformational flexibility and the propensity for phase separation^33,47^.

To quantify the effect of missense mutations on phase separation, we compared the mutant and wild-type proteins using their LLPS propensity scores. We computed per-residue scores by applying our ensemble classifier to 21-residue windows across the mutant and wild-type protein sequences. We defined the mutation-induced change in LLPS propensity as the sum of the per-residue score differences between the mutant and wild-type (WT) proteins, denoted mut(i) and WT(i), respectively.

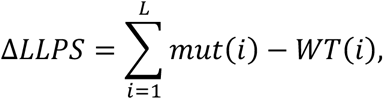

where L is the sequence length. Positive values of ΔLLPS indicate a gain in LLPS propensity, and negative values indicate a loss. To facilitate interpretation of ΔLLPS magnitude, we also reported ΔLLPS z-scores for individual mutations, calculated by centering each score on the mean and scaling by the standard deviation of the overall ΔLLPS distribution. In addition to analyzing naturally occurring variants, we performed in silico mutagenesis to systematically assess the effects of amino acid substitutions across protein sequences. For each protein, we introduced single-amino-acid substitutions at every position, generating all possible missense mutations. We input each mutated sequence and its corresponding structure-aware tokens into the ensemble classifier. This approach enables high-throughput identification of mutations that strongly modulate phase separation.

## Source code and data availability

The training, validation, and test datasets used in this study, along with predicted phase-separation scores for the human proteome, predicted phase-separation scores mapped to cellular localizations, and predicted effects of ClinVar and the denovo-db mutations on phase separation, are provided in the supplementary material. The source code for model training and inference developed in this study is available at https://github.com/kumarlab-compomics/LLPS_Predict.

## Acknowledgments

We thank Alexander Turco for assistance with visualizing the schematic figures presented in this work. We also thank Luke Zhang for valuable discussions.

