## supplementary figures for "Interpretable Prediction of Phase Separation and Disease Variant Effects in Intrinsically Disordered Regions"

### Supplementary Materials

#### Method: Calibration Of Per-Residue Propensity Scores

We used a sliding window method to identify sequence segments that promote phase separation. A fixed-length window of 21 residues moved sequentially along each protein sequence and its corresponding structure-aware tokens. Our ensemble classifier processed each window independently and assigned a propensity score to the central residue of the window. Because sequence length is known to correlate with phase-separation propensity, applying a classifier trained on full-length sequences to short sequence windows introduces a shift in the score distribution. To account for this discrepancy, we calibrated the per-residue scores using Platt scaling. The calibrated propensity score for the i-th residue is defined as:

$$p_i = \frac{1}{1 + \exp\left(Af_i + B\right)}$$

where $f_i$ is the logit produced by the final linear layer of the model for the corresponding sequence segment. $A$ and $B$ are scalar parameters learned from data. These parameters were estimated by fitting the Platt's scaling model on 22 LLPS proteins from the training set with experimentally validated phase-separating regions available in the PhaSePro database. Notably, when $A = -1$ and $B = 0$, this formulation reduces to the sigmoid function used during model training on full-length sequences. For the 21-residue sliding windows, the fitted parameters were $A = -1.01$ and $B = -2.61$. This calibration step adjusts for differences in score distributions between full-length sequences and short windows, enabling consistent interpretation of per-residue scores and the use of a unified cutoff (0.5) for identifying critical regions.

### Model Architectures

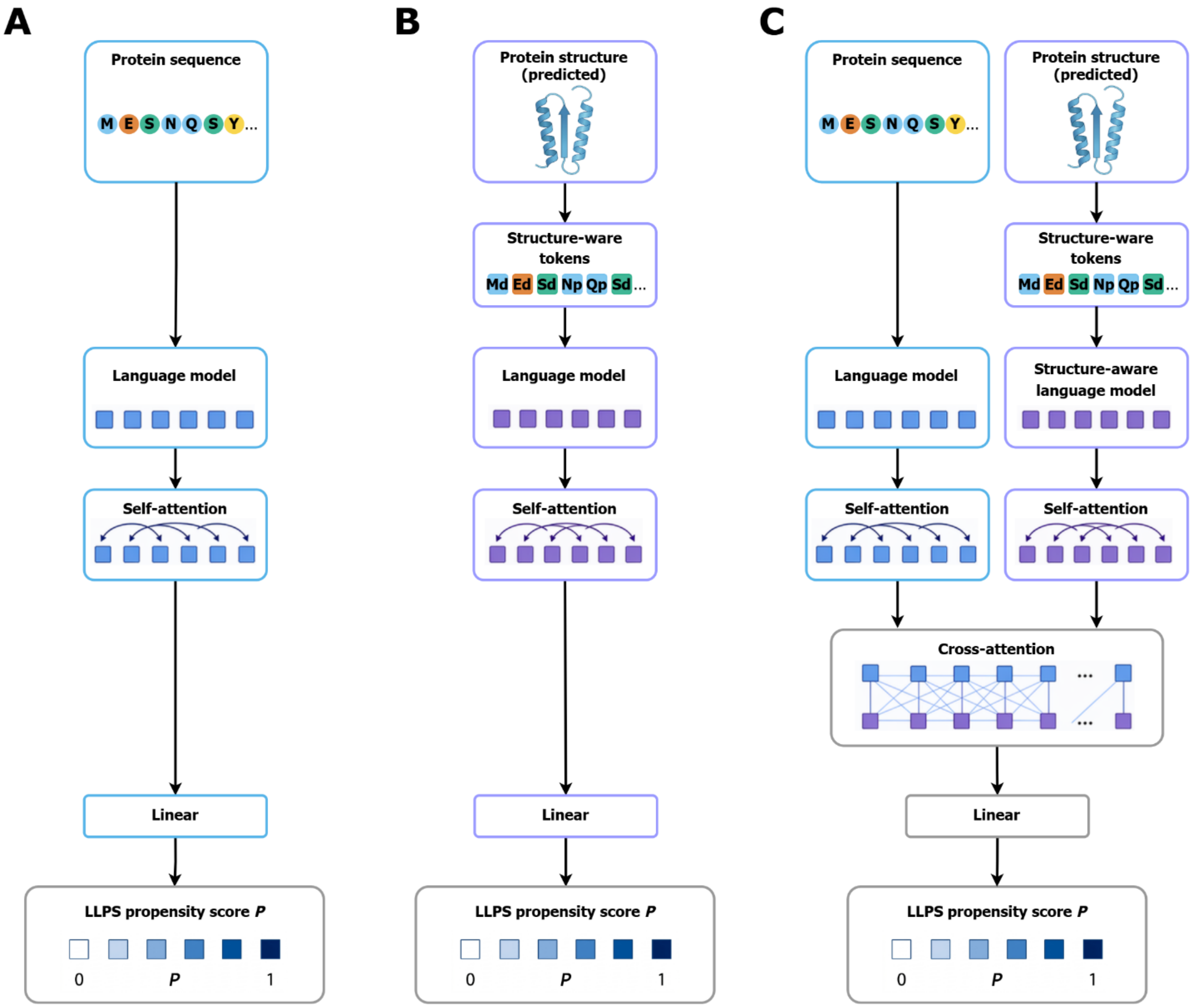


**Figure S1: Architecture of sequence-based, Structure-aware, and ensemble LLPS prediction models.** We first trained two language-model-based predictors and subsequently trained an ensemble model that integrates their learned representations. **A)** Sequence-based model using residue-level embeddings generated by the protein language model ESM2. **B)** Structure-aware model using residue-level embeddings generated by the protein language model SaProt, which incorporates structural information derived from protein three-dimensional structures. **C)** Ensemble model that combines ESM2 and SaProt representations through a multi-head cross-attention layer to leverage complementary sequence and structural information for LLPS prediction.

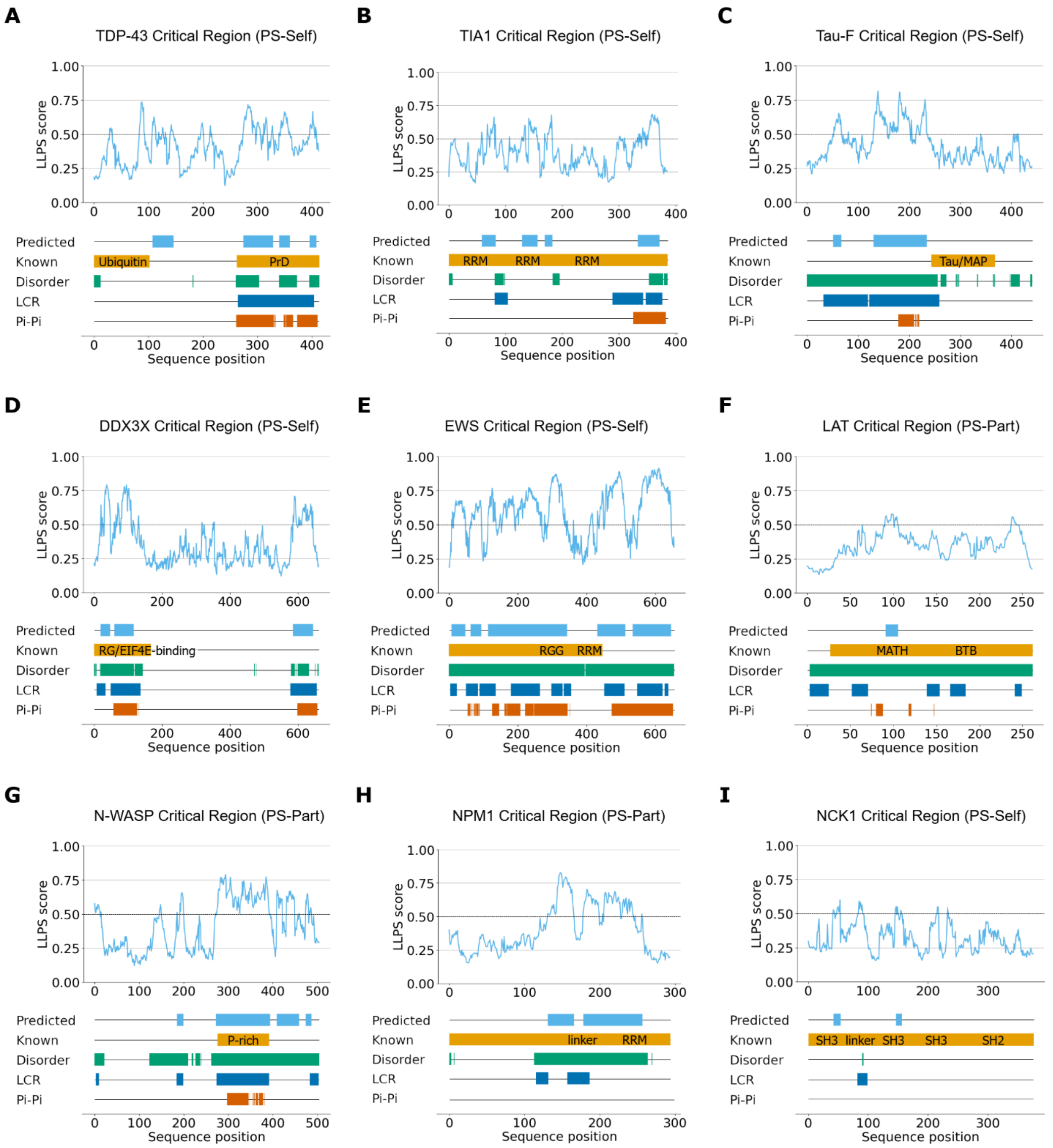


**Figure S2: Critical regions of phase-separating proteins. A-E)** Residue-level LLPS propensity profiles of the self-separating LLPS proteins **A)** TDP-43, **B)** TIA1, **C)** Tau-F, **D)** DDX3X, and **E)** EWS. **F-I)** Residue-level LLPS propensity profiles of the partner-dependent LLPS proteins **F)** LAT, **G)** N-WASP, **H)** NPM1, and **I)** NCK1. From top to bottom, the tracks show predicted critical regions, experimentally identified phase-separating regions, intrinsic disorder, low-complexity regions (LCRs), and predicted π-π interaction propensity. The self-separating and partner-dependent labels were collected from the PhaSePro database.

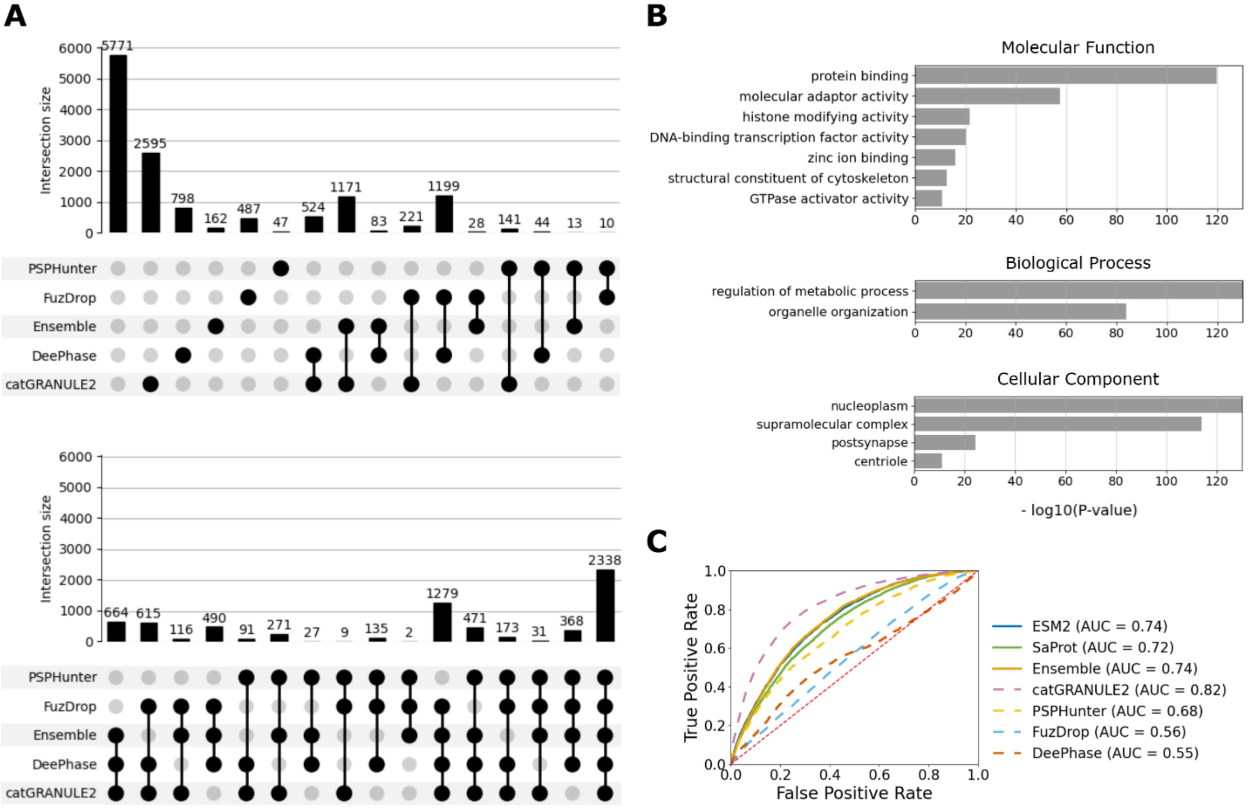


**Figure S3: Human proteome-wide predictions of LLPS proteins by five different methods.** **A)** Numbers of predicted LLPS proteins in the human proteome by five methods, including PSPHunter, FuzDrop, DeePhase, catGRANULE2, and our method (Ensemble). We used the cutoffs suggested by Pintado-Grima et al., 2025, for all methods except ours. **B)** Pathway enrichment analyses performed on the common set of 2,338 proteins identified by all five methods. **C**) Lower-bound AUC performance of different LLPS predictors on the human proteome. All known LLPS proteins were labeled as positives, and the remaining proteins in the proteome were labeled as negatives.

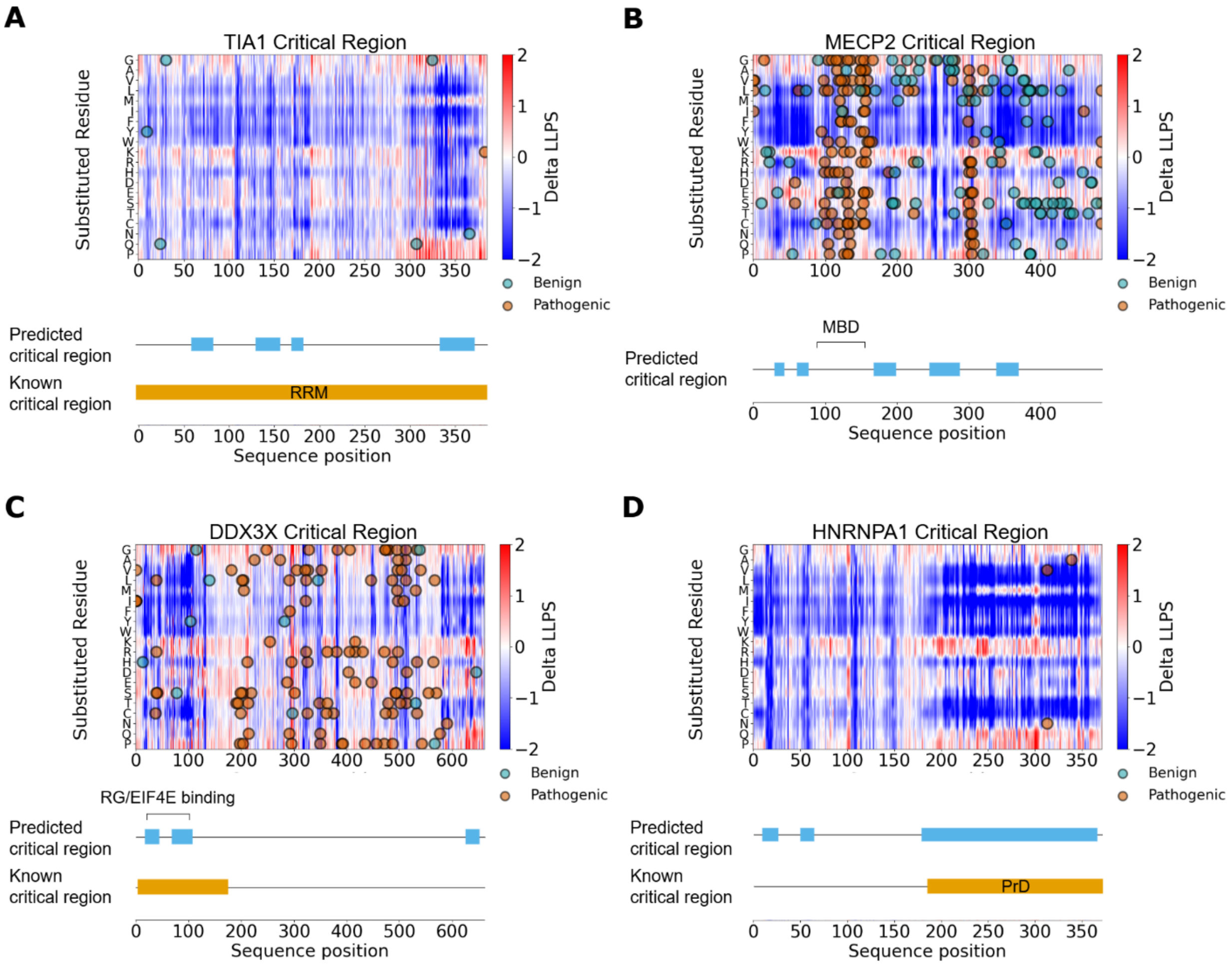


**Figure S4: in silico saturation mutagenesis analyses of phase-separating proteins.** For the known phase-separating proteins TIA1, MECP2, DDX3X, and HNRNPA1, we conducted saturation mutagenesis analyses by replacing each residue with other residues and quantifying the effects of the resulting single-point mutations with the ΔLLPS score. A negative ΔLLPS score indicates a decrease in LLPS propensity, whereas a positive ΔLLPS score indicates an increase. The points overlayed on the mutagenesis heatmaps represent missense mutations from the ClinVar and denovo-db databases. The blue and orange tracks above each mutagenesis heatmap illustrate the predicted and known phase-separating regions respectively.

**<u>Datasets</u>**

| Dataset | Source | Label | Number of Samples | Total |
|---|---|---|---|---|
| Training | Li et al., 2024 | LLPS | 748 | 9,250 |
| | DrLLPS | PS-Self | 52 | |
| | | PS-Part | 3,598 | |
| | LLPSDB | PS-Self | 174 | |
| | | PS-Part | 31 | |
| | PhaSePro | PS-Self | 10 | |
| | | PS-Part | 12 | |
| | PDB | Non-LLPS | 4,625 | |
| Validation | DrLLPS | PS-Self | 55 | 464 |
| | | PS-Part | 116 | |
| | LLPSDB | PS-Self | 56 | |
| | PhaSePro | PS-Self | 5 | |
| | PDB | Non-LLPS | 232 | |
| Test | Pintado-Grima et al., 2025 | PS-Self | 285 | 2,409 |
| | | PS-Part | 222 | |
| | | PS-Self & PS-Part | 42 | |
| | | Structured Non-LLPS | 930 | |
| | | Disordered Non-LLPS | 930 | |
| Cellular Localization | Hardenberg et al., 2020 | Cellular location | 15,283 | 15,283 |
| Pathway Enrichment | PDB | Predicted PS-Self | 1,050 | 20,374 |
| | | Predicted PS-Part | 1,664 | |
| | | Other | 17,981 | |
| Mutations | ClinVar & denovo-db | Predicted PS-Self | 1,539 | 11,089 |
| | | Predicted PS-Part | 3,541 | |
| | | Predicted Non-LLPS | 6,009 | |

**Table 1: Datasets used to train, validate, test, and evaluate our models.** We removed sequences shorter than 50 residues and used CD-HIT to reduce sequence redundancy in the training, validation, and test sets. Pairwise sequence similarity within the validation set was limited to below 50% (cd-hit -c 0.5 -n 2). We ensured that all sequences in the validation and external test sets share below 50% identity with any sequence in the training set (cd-hit-2d -c 0.5 -n 2). All reported sample counts correspond to the datasets after applying these filters.

**SaProt Structure-Aware Tokens**

| SaProt structural placeholder | PS-Self vs. non-LLPS AUC | F1 | PS-Part vs. non-LLPS AUC | F1 |
|---|---|---|---|---|
| Placeholder pLDDT < 90 | 0.93 | 0.83 | 0.92 | 0.79 |
| Placeholder pLDDT < 70 | **0.94** | 0.83 | **0.93** | **0.80** |
| Placeholder pLDDT < 50 | **0.94** | **0.84** | **0.93** | **0.80** |
| No placeholder | **0.94** | **0.84** | **0.93** | **0.80** |

**Table 2: Structure-aware language model input ablation study.** The SaProt model requires structure-aware tokens, in which protein structures are encoded as tokens using the 3Di alphabet introduced by Foldseek. In this alphabet, each amino acid (denoted by the upper-case letters) is paired with a discrete structural token (denoted by the lower-case letters). For regions with low-confidence AlphaFold2 predictions, the structural component can be replaced by a placeholder ("#"). A commonly used threshold for assigning placeholders is a predicted local distance difference test (pLDDT) score below 0.7. We performed an ablation experiment by training the ensemble model with different pLDDT thresholds for placeholder assignment. Based on validation set performance, increasing the number of placeholders led to a slight decrease in model accuracy in both PS-Self and PS-Part classification. For example, for PS-Self vs. non-LLPS classification, the ensemble model trained without placeholders achieved an AUC of 0.94, whereas introducing placeholders for residues with pLDDT < 90 resulted in a reduced AUC of 0.93. Similarly, for PS-Part vs. non-LLPS classification, the AUC decreased from 0.93 to 0.92 under the same threshold. Retaining all structural tokens without introducing placeholders yielded the best performance. Therefore, we used the full set of structure-aware tokens without placeholders in our final models.

**Classification Model Cutoffs**

| PS Model Cutoffs | False Positive Rate | Sensitivity | F1 | Matthews Correlation Coefficient |
|---|---|---|---|---|
| P = 0.2 | 0.34 | 0.94 | 0.82 | 0.62 |
| P = 0.3 | 0.27 | 0.93 | 0.85 | 0.67 |
| P = 0.4 | 0.21 | 0.90 | 0.85 | 0.69 |
| P = 0.5 | 0.15 | **0.88** | **0.87** | **0.73** |
| P = 0.6 | 0.13 | 0.85 | 0.86 | 0.72 |
| P = 0.7 | **0.11** | 0.80 | 0.84 | 0.69 |

**Table 3: Ensemble model performance using different cutoffs.** We measured the performance of the ensemble model on the validation set (LLPS vs. non-LLPS). All proteins with predicted scores above the cutoff are classified as phase-separating proteins. Among all cutoffs tested, the 0.5 cutoff resulted in the highest F1 score and Matthews Correlation Coefficient, so we used this cutoff for all downstream analyses.
